# The dominance of coinfecting parasites’ indirect effects on host traits

**DOI:** 10.1101/2023.02.12.528182

**Authors:** Daniel I. Bolnick, Sophia Arruda, Christian Polania, Lauren Simonse, Arshad Padhiar, Andrea Roth, Maria L. Rodgers

**Affiliations:** Department of Ecology and Evolutionary Biology, University of Connecticut, Storrs CT 06269, USA; Department of Biological Sciences, North Carolina State University, Morehead City NC 28557, USA

**Keywords:** Coinfection, fibrosis, ecological immunology, *Gasterosteus aculeatus*, gene flow, effective migration rate, *Schistocephalus solidus*, threespine stickleback

## Abstract

Indirect genetic effects (IGEs) exist when there is heritable variation in one species’ ability to alter a second species’ traits. For example, parasites can evolve disparate strategies to manipulate host immune response, whether by evading detection or suppressing immunity. A complication arises during coinfection, when two or more parasite genotypes may try to impose distinct IGEs on the same host trait: which parasite’s IGE will be dominant? Here, we apply the notion of dominance to IGEs during coinfection. Using a mathematical model we show that the dominance of IGEs can alter the evolutionary dynamics of parasites. We consider a resident parasite population receiving rare immigrants with a different immune manipulation trait. These immigrants’ relative fitness depends on resident prevalence (e.g., the probability immigrants are alone in a host, or coinfecting with a native), and the dominance of the immigrant’s IGE on host immunity. Next, we show experimentally that the cestode *Schistocephalus solidus* exerts an IGE on a host immune trait: parasite antigens from different populations produced different intensities of fibrosis. We then evaluated IGE dominance, finding evidence for overdominance (coinjected antigens induced an even stronger host immune response) which would be detrimental to immigrants when resident prevalence is high. This combination of experimental and modeling results shows that parasites do exhibit IGEs on host traits, and that the dominance of these IGEs during coinfection can substantially alter parasite evolution.

## INTRODUCTION

Coinfection is the typical state in natural populations (Poulin 2007; Graham 2008; Seabloom et al. 2015; Diuk-Wasser et al. 2016; Marchetto and Power 2018; Bolnick et al. 2020). Most individual animals are infected by multiple parasite species, as well as by multiple individuals of a given parasite species (e.g., Fig. S1). The consequences of such coinfections can include changes in parasite growth rates (Lass et al. 2013), host traits (Mabbott 2018), duration of infection (Krause et al. 1996; Diuk-Wasser et al. 2016), and disease severity (Krause et al. 1996; Graham et al. 2005; Gibson et al. 2011). Coinfecting parasites may compete, mutually reducing their fitness (Blackwell et al. 2013). Alternatively, coinfection may be mutualistic, facilitating both parasites’ survival and virulence (West and Buckling 2003). These parasite-parasite interactions can arise from (1) direct molecular interference (Damian 1997; Ezenwa et al. 2010; Harnett 2014), (2) competition for shared host resources (Budischak et al. 2018; Wedekind and Rüetschi n.d.), or (3) indirectly via changes in host immune responses (Ezenwa et al. 2010; Mabbott 2018; Ling et al. 2020).

Indirect interactions between parasites arise because one or both coinfecting species alter host traits, which in turn affect the fitness of either parasite. These host trait changes are an example of an ‘indirect genetic effect’ (IGE, (De Lisle et al. 2022)), or an ‘extended phenotype’ (Geffre et al. 2017). That is, genetic variation among parasites can induce phenotypic variation among hosts. Most notably, parasites are well known to suppress, misdirect, or otherwise manipulate host immune traits (Damian 1997; Schmid-Hempel 2008; Geffre et al. 2017; Mabbott 2018; Chulanetra and Chaicumpa 2021). Immune evasion may entail simple molecular camouflage, like the evolution of protein antigens that hosts fail to detect. Examples include antigen sequences that evade recognition by Major Histocompatibility Complex (MHC) proteins (Hunt et al. 1992; Cnops et al. 2015), or mimic host self-antigens that the immune system ignores (Revilleza et al. 2011; Miller et al. 2019). Alternatively, parasites may evolve strategies to actively interfere with a host’s immune defenses via molecular signals in their excretory-secretory products (ESPs), also known as the secretome (Hiller et al. 2004; Hotterbeekx et al. 2021; Wititkornkul et al. 2021). This mixture of molecules that a parasite releases can disrupt a host’s physiological functions to suppress immune function (Hewitson et al. 2009; Harnett 2014), or to misdirect it into an ineffective response (Sisquella et al. 2017).

Coinfection generates the potential for conflict between two (or more) parasites’ extended phenotypes (Mabbott 2018). For example, upon reaching a threshold size, the tapeworm *Schistocephalus solidus* induces behavioral changes in its fish host to facilitate bird predation (Piecyk et al. 2019; Berger et al. 2021). Coinfection between a large and small tapeworm generates a conflict in which the immature parasite manipulates the host to be cautious, while the larger mature parasite induces risky behavior (Barber and Huntingford 1995; Hafer and Milinski 2015, 2016). When such conflicts arise, a key question is which trait does the host exhibit? Is one parasite’s indirect genetic effect dominant, and another recessive? At present, little is known about the dominance of parasites’ IGEs during coinfection, or the consequences of such dominance. Here, we first present a mathematical model showing that the IGE dominance of coinfecting parasites can affect the relative fitness of parasite genotypes, and thereby alter parasites’ evolutionary dynamics. Second, we provide experimental data showing that a parasitic tapeworm *Schistocephalus solidus* exerts indirect genetic effects on their host (threespine stickleback): some tapeworm genotypes induce a stronger host immune response (fibrosis) than others. We then use a coinfection assay to show that the stronger IGE is overdominant: coinfection causes a stronger immune response than either parasite alone, which will inhibit gene flow between parasite populations and thereby alter the potential for host and parasite local coevolution.

### Immunological dominance in coinfection

Consider the case of coinfection between two parasite genotypes, which exert different indirect genetic effects (IGEs) on the host (De Lisle et al. 2022). One genotype (RR) induces a strong immune response, and the other genotype (rr) does not (Fig. 1). If they coinfect, does the host initiate a strong immune response (responding to RR), or not (responding to rr)? Biologically, this will depend on the mechanism of their IGEs.

**Figure 1:**
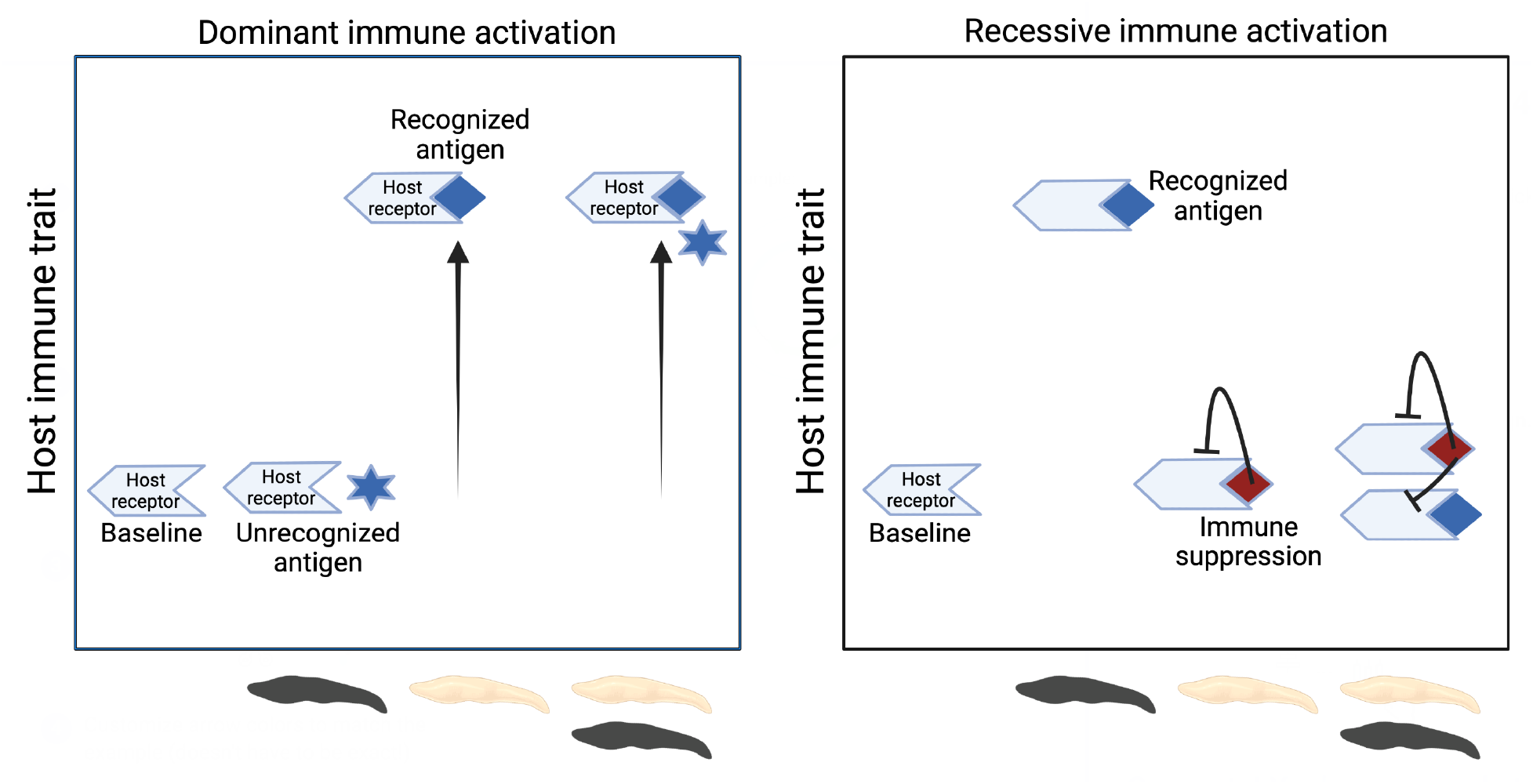
The immunological effect of coinfection can be thought of as a case of genetic dominance, as both parasites exert extended phenotypic effects on host traits. Here we illustrate two scenarios, showing a host immune trait in the absence of a parasite (‘baseline’), and with either of two parasite genotypes (indicated by white, or black tapeworms) individually or together. A) Dominant immune activation: the host activates a stronger immune trait when its receptor successfully recognizes the second parasite antigen (diamond symbol), whether or not an unrecognized antigen (star) is present. B) Recessive immune activation: the host receptor recognizes both parasite antigens and would initiate a strong immune response. However, the second parasite antigen (red diamond) has an immune suppressive effect. As a result, the immune response is low in the coinfection. Alternatively, the coinfection might induce an intermediate host response (akin to additive genetic effect), or perhaps induce a more extreme trait (‘overdominance’ or ‘underdominance’).

One possibility is that the host recognizes an RR-parasite antigen and initiates a response (e.g., because its MHC antigen-binding groove binds to the parasite protein), but does not respond to rr because the host fails to recognize the slightly different rr antigen. The coinfection (RR/rr) will be recognized by the host because of the positive presence of the R antigen, and a strong immune response will ensue. In effect, the R allele’s IGE is dominant. A second scenario could be that the host immune system recognizes both RR and rr parasite genotypes, but the rr genotype secretes an immune suppressive product inhibiting an immune response. In a coinfection (RR/rr), the immune suppressive product is present and host immunity inhibited, so the r allele has the dominant IGE. Of course, these two possibilities are not mutually exclusive and intermediate outcomes are conceivable. The host might be better at recognizing R and be actively suppressed by r, which might result in an intermediate phenotype for coinfections.

The full range of possible coinfection outcomes can be encapsulated by applying the classic quantitative genetics view of dominance to the notion of IGE (Fig. 1). We can view the RR/rr coinfection as if it were a heterozygote, and estimate a dominance coefficient, *d.* When the coinfection induces the weaker immune response (rr’s IGE) then *d* = 0. Conversely when coinfection induces the stronger immune response (RR’s IGE), *d* = 1.0. If the effects are intermediate, 0 < *d* < 1 (exactly additive when *d* = 0.5). Transgressive variation is possible as well (e.g, overdominance when *d* > 1.0). The dominance of coinfection IGEs is not currently known, empirically. Nor have evolutionary models of IGEs (e.g., (De Lisle et al. 2022) considered the impact of coinfection on host-parasite coevolution, local adaptation, or epidemiological dynamics. In this paper we first develop a mathematical model to demonstrate that IGE dominance matters for the relative fitness of parasite genotypes (and hence, parasite evolution). Then, we present an empirical estimate of IGE dominance.

## MODEL: THE EFFECTS OF IGE DOMINANCE ON IMMIGRANT FITNESS

### Model framework

Here we consider the short-term effects of immunological dominance of IGEs on the relative fitness of immigrant versus resident parasites. Our model focuses on a single geographically bounded population of hosts with a native population of parasites, that receives occasional immigration of parasites from other such host populations. When immigrants’ fitness is greater than that of residents, the immigrant genotype will establish and increase in frequency (at least in the short term; over long time-scales co-evolution and frequency-dependent interactions would require a more extensive analysis). When immigrants fitness is less than that of residents, selection reduces the effective immigration rate and will tend to maintain genetic differences between parasite populations. This reduced gene flow may facilitate parasite local adaptation, although if the hosts coevolve with the parasite then reduced parasite gene flow may facilitate host local adaptation instead (Gandon and Michalakis 2002; Hoeksema and Forde 2008).

The infection intensity of resident or immigrant parasites (I_r_ and I_i_ respectively) depend on their infection rates λ_r_ and λ_i_ following a Poisson distribution:

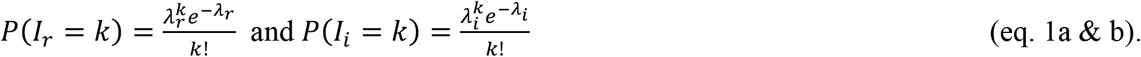

Note that we assume that, by definition, resident infection rates exceed the rate of immigrant infections (λ_r_ >> λ_i_). The probability a given host is infected by residents, or by immigrants, (a.k.a. infection prevalence) is therefore:

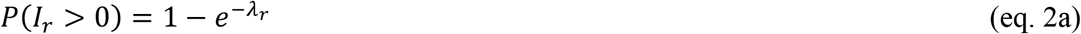

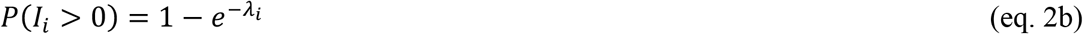

From this we can calculate the proportion of hosts that are infected only by residents, only by immigrants, or coinfected:

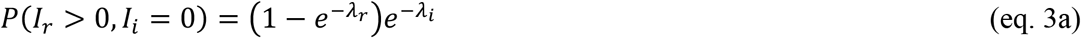

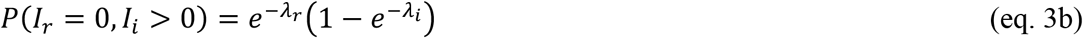

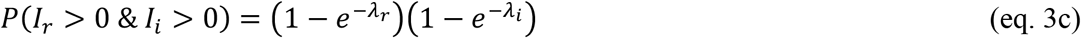

The expected total infection intensity (*T* = I_r_ + I_i_; residents and parasites) is:

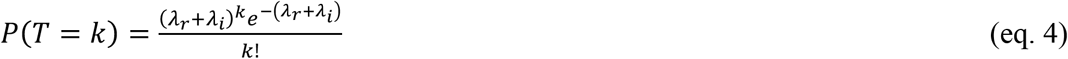

Having defined the relative rate of coinfections, and total intensity (affecting within-host competition), we can define parasite fitness, which we split into three multiplicative components. First, each genotype has a baseline fitness unaffected by crowding or host immunity, *ω_ir_* and *ω_ii_*. Assuming there is some parasite local adaptation, *ω_ii_* < *ω_ir_* = 1. Second, both parasite genotypes are harmed by parasite-parasite competition within a host, so that baseline fitness is multiplied by (1 — *αT*). 1/*α* represents the coinfection carrying capacity *K*, the density at which overcrowding reduces parasite fitness to zero. Third, each host has a probability *γ* of initiating an immune response, which kills all parasites present.

Indirect genetic effects (IGEs) arise because parasite genotypes induce different host immune responses *γ*. We assume that hosts are at least in part locally adapted, evolving a stronger immune response to native than to immigrant parasites (*γ*_r_ > *γ*_i_). Phrased another way, resident parasites have larger IGE on host immune response. Coinfected hosts mount an immune response with probability gc, which depends on the IGE dominance

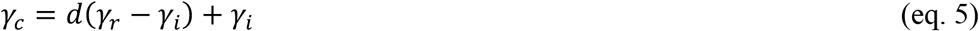

If hosts fail to detect immigrant parasites, then we expect γ_i_ < γ_c_ = γ_r_. so *d = 1*. Alternatively if immigrants suppress host immunity then γ_i_ = γ_c_ < γ_r_. More realistically coinfection may represent some intermediate immune response (γ_i_ < γ_c_ < γ_r_) so 0 < *d* < 1. We thus have different fitnesses for resident and immigrants, without coinfection:

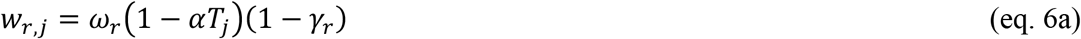

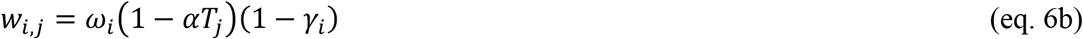

and with coinfection:

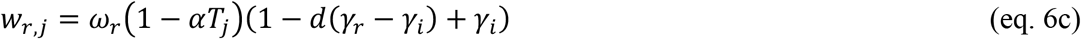

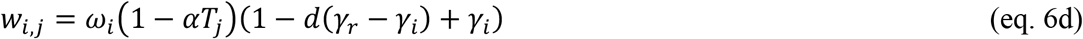

To obtain average resident and immigrant fitness we then average these across the range of possible infection intensities (the distribution of T which affects competition). The competition term (1 – *αT_j_*) becomes 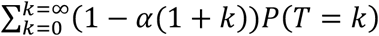. The (1+k) term lets us condition on the focal parasite being present. This yields a simple expectation for the competition term, 1 – *α* – α(*λ_r_* + *λ_i_*). Next, we average over the frequencies of single and coinfections, resulting in the expected resident fitness

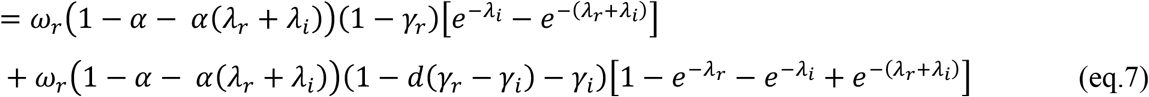

And the immigrant fitness is

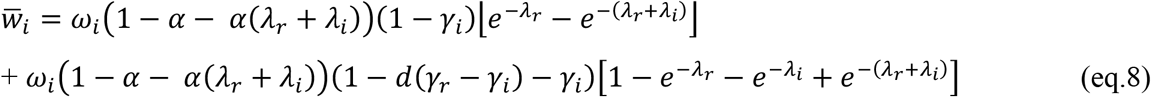

The immigrant’s invasion fitness is 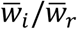, which determines how immigrant success will depend on infection rates (dominated by resident abundance *λ_r_*), and immune dominance *d.* We calculated numerical solutions to equations 7 and 8, iterating through a range of values of IGE dominance (d from 0 to 2 in steps of 0.1), and resident infection loads *λ_r_* varying from 0.1 to 5. For all simulations we kept immigrants rare (*λ_i_* = 0.01), imposed weak costs of crowding (*α* = 0.02, implying a maximum carrying capacity of T=50 parasites per host). These parameters are approximately realistic reflections of ranges observed in *S.solidus* infections of stickleback. We assume a slight advantage of residents in density-independent local adaptation (*ω_i_* = 0.95), and a strong host immune response to residents but no immune response to immigrants (*γ_r_* = 0.5, *γ_i_* = 0).

### Model results

Our analyses show that the dominance of parasite indirect genetic effects alters the relative fitness of immigrant versus resident parasites, but this effect depends on resident parasite abundance (Fig. 2). We first consider the case where immigrants are simply not recognized by hosts, which is reasonable because immigrants are rare and impose little if any selection on hosts’ pattern recognition molecules. In this case, coinfection with residents should induce a typical immune response against residents (d = 1, Fig. 1A) to the detriment of both resident and coinfecting immigrant parasites. As a result, when resident infection is common (e.g., *λ_r_* = 2), residents and immigrants are about equally fit, both being substantially harmed by host immunity (even though immigrants would not induce immunity on their own). However, when resident abundance is low (e.g., *λ_r_* = 0.1), then immigrants attain a higher mean fitness than residents because most immigrants are alone and escape immune detection, unlike the residents.

**Figure 2.**
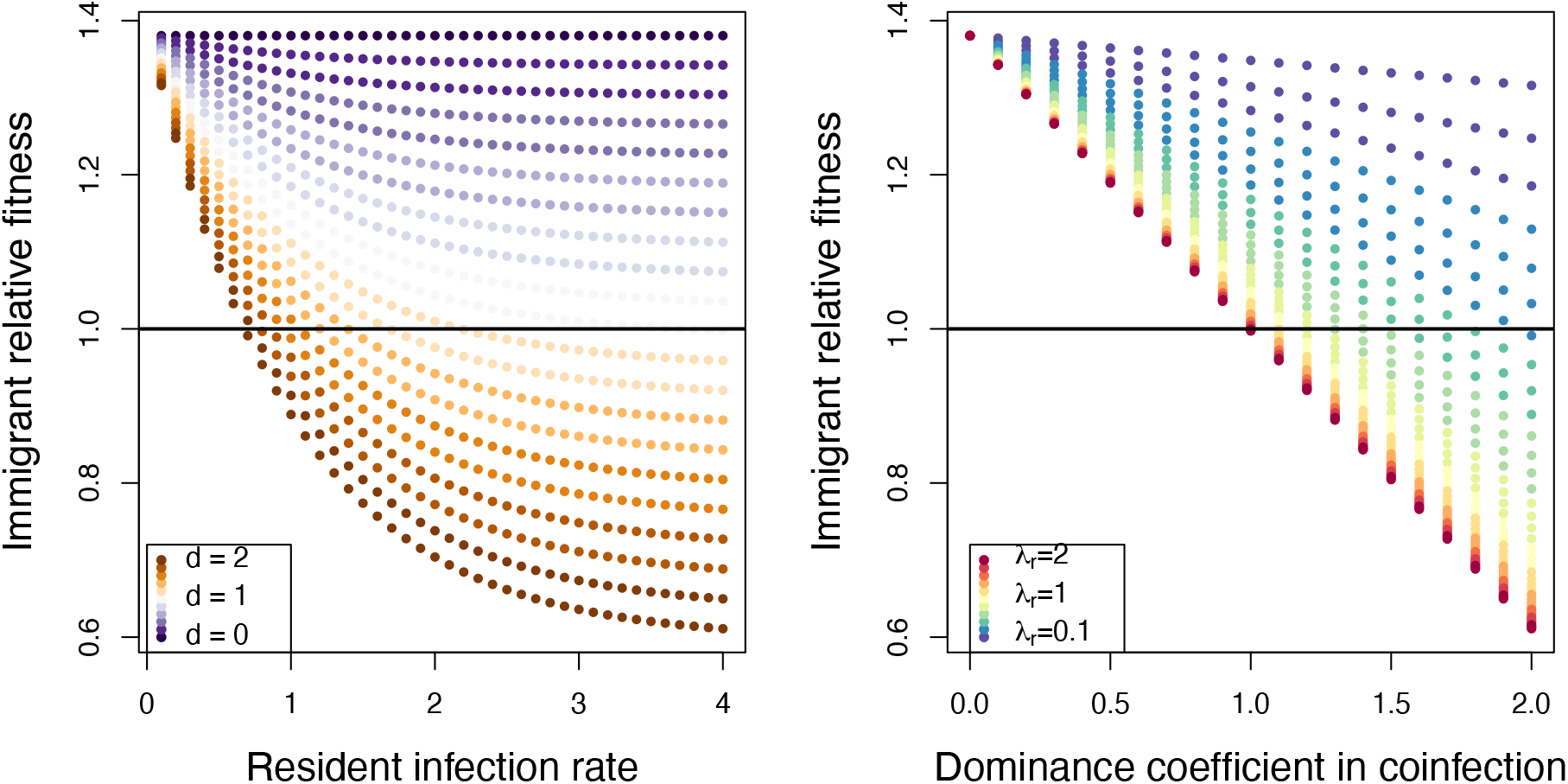
The relative fitness of an immigrant parasite genotype, compared to resident parasite fitness 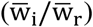 depends on the interaction between immunological dominance (d) and resident infection rates (λ_r_). The two panels present the same data, just focusing on different parameters as the x axis.

Alternatively, immigrants might suppress host immune function (Fig. 1B), for instance if hosts lack tools to counteract immunosuppression by foreign parasites. Coinfected hosts would then fail to respond to infection (d = 0). This recessive immune IGE gives immigrants a substantial benefit over residents due to their safety from host immune attack regardless of coinfection. However, when coinfection is common then the immigrants’ immune suppression rescues their resident competitors who would have otherwise been attacked by the host immune system. Thus, when d = 0, immigrants outperform residents but this benefit of immune suppression is eroded by increased resident abundance.

A third possibility is that coinfection by two genotypes causes a higher immune response than is seen with either parasite genotype alone. Such overdominance (d > 1) reduces immigrant fitness, especially when coinfection is highly likely. At low resident abundance, immigrants are still able to escape host immunity and attain higher fitness than residents (in the parameter space considered here, the benefit of immune evasion exceeds resident parasites’ baseline advantage from local adaptation). The resident advantage is reduced weakly by immune dominance, but because coinfection is rare this has little effect. However, when coinfection is common, immune dominance can entirely reverse the relative fitness of residents and immigrants: the former are more fit when there is overdominance and frequent infection, the latter more fit when coinfection is rare. This result demonstrates that the effective rate of gene flow into a resident parasite population will tend to be reduced by host immune over-dominance, especially when resident infection rates are high. Immune dominance thus modulates parasite gene flow in a density-dependent manner. For brevity, we do not explore the long-term evolutionary dynamics arising from these effective migration rates. However, the fact that IGE dominance changes immigrants’ invasion fitness strongly suggests that it will modulate the extent of genetic divergence among parasite populations. Because the effective rate of gene flow affects the parasites’ capacity to establish local adaptation (to the host, or to abiotic conditions), we consider this invasion fitness analysis to be strong evidence that IGE dominance has evolutionary effects. Further analyses of long-term dynamics are warranted, but will certainly entail a mix of density- and frequency-dependent effects. In this regard, these results add to a growing body of work suggesting that selection for or against immigrants can be both density- and frequency-dependent (Bolnick and Stutz 2017).

## EXPERIMENTAL METHODS

### Model system

We conducted immune challenge experiments to evaluate aspects of coinfecting immune dominance, using threespine stickleback *(Gasterosteus aculeatus)* as a study system. Stickleback are commonly infected by a tapeworm *Schistocephalus solidus* which can grow to over half the fishes’ body mass (Weber et al. 2017). An individual parasite is acquired when stickleback consume an infected copepod, and the tapeworm passes through the intestinal wall to grow to mature size in the fish’s body cavity. While infecting its fish host, *S.solidus* secretes compounds that manipulate its host immune response (Scharsack et al. 2004, 2007a, 2013; Berger et al. 2021) and altering behavior to increase susceptibility to bird predation (Barber and Huntingford 1995; Talarico et al. 2017). The parasite mates in the birds’ intestines before eggs are defecated - possibly into a different water body (Shim et al. 2022b). As a result of this life history the tapeworm has a higher dispersal rate and far more gene flow than their host (Shim et al. 2022b).

Despite this high dispersal capacity, infection rates vary between stickleback populations, and may be absent or vanishingly rare in marine populations and some lakes, but infect a majority of individuals in other geographically nearby lakes (Weber et al. 2017). In lakes with low infection prevalence, most infected fish harbor only a single tapeworm (coinfection is rare), but in lakes with high prevalence up to a third of individuals may be infected by more than one worm (Fig. S1).

Host ecology and immune genotype both contribute to this geographic variation in *S.solidus* infection prevalence and intensity. Some stickleback populations consume more copepods (the parasites’ first host) than other populations, and some populations are genetically predisposed to resist infection (Weber et al. 2017, 2022). In particular, stickleback from a subset of lakes can mount an effective immune response to the tapeworm (Weber et al. 2022). Exposure to the parasite induces fibroblast activation to generate extensive fibrosis, scar-tissue lesions forming a cocoon around the organs and parasite, sometimes binding the organs tightly to the body wall. This fibrosis reduces parasite growth dramatically, and sometimes allows the fish to encase the parasite in a cyst leading to parasite death (Weber et al. 2022). This fibrosis response is heritable, differs between populations, and can be induced through experimental vaccinations of either alum (an immune adjuvant) or *S.solidus* protein extract (Hund et al. 2022).

Experimental infections by *S.solidus* often have low success rates (e.g., 10 – 20%) due to host immunity (Weber et al. 2017), so live coinfection experiments are inefficient. On average nearly 100 fish would need to be experimentally co-exposed, to generate a few successful coinfections. In contrast, injecting cestode protein reliably induces extensive changes in host fibrosis and gene expression (Hund et al. 2022), providing a practical alternative to live coinfection. We used this vaccination methodology to evaluate two related questions. In Experiment 1, we asked whether tapeworm proteins obtained from different source populations of parasites vary in their propensity to induce host fibrosis (e.g., is there variation in their indirect genetic effects?). Experiment 2 then evaluated whether this variance in fibrosis exhibits dominance. By coinjecting proteins from two tapeworm genotypes (one inducing fibrosis, the other not), we estimated the dominance of their IGEs. Do coinjected hosts exhibit fibrosis comparable to the pro-fibrotic parasite genotype, the non-fibrotic genotype, intermediate, or transgressive?

### Experiment 1: Parasite indirect effects on host immune traits

In June 2019, we collected threespine stickleback from Roselle Lake (50°31’13”N, 126°59’12”) using minnow traps. Previous work revealed that stickleback from this lake, when injected with alum or cestode protein, induce rapid fibrosis (Hund et al. 2022). Using standard in-vitro fertilization methods for stickleback, we bred the wild-caught fish and transported the fertilized eggs back to the lab. These fish were raised for two years at the University of Connecticut. Field collections were conducted with approval from the British Columbia Ministry of Forests, Lands, Natural Resource Operations and Rural Development (Fish Collection Permit NA19-457335).

Roselle Lake stickleback were injected with cestode protein extracts, each fish receiving protein obtained from one of four parasite populations. Tapeworms were dissected from wild-caught stickleback from three lakes in British Columbia (Roselle Lake; Boot Lake, 50°03’12”N, 125°31’47”W; Gosling Lake, 50°03’47”N, 125°30’07”W), and Cheney Lake in Alaska (61°12’06”N, 149°45’37”W). Field-trapped infected fish were frozen and shipped back to the lab, then thawed and dissected to acquire parasites. Individual parasites were sonicated in phosphate-buffered saline (PBS) on ice, and the resulting suspension centrifuged at 4000 rt/min at 4 °C for 20 minutes. Using the clear, upper fraction of the resulting solution, we assessed protein concentration in triplicate using RED 660™ protein assay (G-Biosciences). Following earlier tapeworm homogenate injection experiments (Vrtílek and Bolnick 2021; Hund et al. 2022), we diluted each sample to 0.45 mg/mL using PBS. All solutions were prepared in a sterile culture hood. Since the efficacy of tapeworm homogenate in stimulating the peritoneal fibrosis response in this population of stickleback had previously been established (Hund et al. 2022), sham injections and antigen-free controls were omitted to focus on between-population comparisons.

We filled syringes aseptically under laminar flow cabinet on the same day as injection and stored them at 4 °C or on ice until used. Before treatment, fish were briefly anesthetized using neutral-buffered MS-222 (~80 mg/L). We injected 20 μL of tapeworm homogenate into the lower left side of the peritoneal cavity of each fish using Ultra-Fine insulin syringes (BD 31G 8mm). During injections, fish were placed on a soaked sponge. Head and gills were covered with a moist paper towel to ensure the fish were adequately hydrated when out of water. Fish were allowed to recover from anesthesia in highly aerated water, then returned to their original tank. Individuals from different treatment groups were housed together to control for tank effects, distinguished by subcutaneous marks of different-colored Visible Implant Elastomer (Northwest Marine Technologies). VIE was injected into dorsal muscle just posterior to the neurocranium. All aspects of the experiment were approved in advance by the University of Connecticut Institutional Animal Care and Use Committee (IACUC Protocol A18-008). Fish that died following injection (an atypical outcome most likely reflecting researcher error damaging an organ) were replaced by new fish given the same treatment, to achieve the target sample size (18 fish per treatment).

Ten days after injection, fish were dissected and peritoneal fibrosis was scored visually along an ordinal scale as in past experiments (Vrtílek and Bolnick 2021; Hund et al. 2022). Scores are: 0 (no fibrosis: organs move freely), 1 (mild fibrosis: light connection of fibrotic threads between liver an intestine, or intestine and swim bladder), 2 (moderate fibrosis: organs difficult to pull apart), 3 (severe fibrosis: organs adhered to peritoneal wall), 4 (very severe fibrosis, adhesion between organs and peritoneal wall is so strong the peritoneum tears when the body cavity is forcibly opened) (see video: https://www.youtube.com/watch?v=yKvcRVCSpWI). Fibrosis was scored by one individual (CMP) who was blind to treatment. Fish mass and sex were recorded, along with elastomer dye marker color to subsequently record the experimental treatment.

We took three approaches to test for fibrosis differences between antigen treatments, to ensure robust inferences. First, we used an ANOVA testing for an effect of parasite source population (4 levels, fixed effect, type II Sums of Squares). Second, we used a Kruskal-Wallis nonparametric test in recognition of the non-normal distribution of the integer ordinal scoring of fibrosis. Third, we used a Bayesian linear model with the R package *rethinking* (McElreath 2016) estimating the overall mean fibrosis score, and treatment-specific deviations from this mean. Specifically we fit a model in which the observed fibrosis values *y_i_* are normally distributed N(ŷ,σ) where the mean differs between treatments such that *y* = *α* + ∑_*i*_ *β_i_I_i_* where β_i_ is the deviation from the mean introduced by antigen genotype *i*, and *I*_i_ is an indicator variable denoting the presence or absence (1 / 0) of the antigen genotype (Roselle, Boot, Gosling, or Cheney). Priors for *β_i_* were normally distributed with mean 0 and standard deviation of 3, the prior for σ was uniform [0,4]. We estimated the mean and 95% credibility interval of the posterior distributions of each parameter. The posterior probability distributions from this analysis served as priors for the second experiment analyses.

### Experiment 2: Dominance of indirect genetic effects during coinfection

As described in detail in the Results (below), Experiment 1 showed that Roselle Lake fish initiate stronger fibrosis when injected with sympatric Roselle Lake tapeworm protein, compared to protein from foreign parasites (Boot or Gosling Lakes, ~115 km away; Cheney Lake, ~1850 km away). In experiment 2 we used coinjection to estimate the dominance of the parasites’ IGE. We injected protein from Roselle tapeworms (pro-fibrotic), Boot tapeworms (non-fibrotic), or Cheney tapeworms (non-fibrotic), alone or in pairwise combination (Roselle + Boot, Roselle + Cheney), with a target of 33 fish per treatment (Table S1), though actual numbers were sometimes slightly lower due to mortality after handling. Treatments were mixed in tanks, distinguished using different elastomer dyes injected subcutaneously. Injections were done in six batches, with each treatment represented in each batch. Across all injection rounds, there was a mortality rate of 2.1% (6 out of 286 fish). Fish were euthanized and dissected 10 days post-injection. Fibrosis was scored as described above, except that two or three individuals scored each fish (one through a binocular microscope, the others watching a live video which was also recorded). The replicate measures were averaged. Due to difficulty in reading some elastomer tags (or, tag loss after injection), 41 fish of uncertain treatment were removed from the dataset prior to analysis.

Because immune responses can be dose-dependent, we tried both additive and substitutive designs for coinjection treatment. Single-parasite injections were 20 μL of 1 mg/ml protein in 0.9x PBS. In the additive design, coinjected fish received 10 μL of 2 mg/ml of protein from each of two parasite genotypes. In this way, the protein mass of each parasite matched their single-injection mass, but the total mass (of both parasites) was higher. In the substitutive design, coinjected fish received 10 μL of 1 mg/ml from each parasite. This matches the total mass of injected protein to the single-parasite treatment, but halves the amount of each protein. A control group of fish received 1x phosphate buffered saline (PBS), which typically does not induce fibrosis. We used an unequal variance t-test to evaluate whether protein concentration (additive versus substitutive) effects subsequent fibrosis. Because we found no significant effect of concentration (later confirmed with a second experiment, Fig. S3), we merged concentrations in the subsequent statistical analyses.

To test for differences in fibrosis between injection treatments we first used linear regression to fit the following linear model:

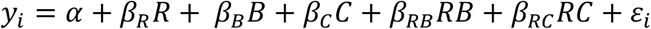

where *y_i_* is the average fibrosis score; *α* is the average fibrosis induced by the control; *R, B,* and *C* are Boolean variables that indicate the presence or absence of Roselle, Boot, and Cheney, respectively; *β_x_* is the effect size of injection type *x* (including interaction effects between RxB or RxC), and *ε_i_* is any random variance. We also used a nonparametric Kruskal-Wallis rank sum test to ensure the results were robust to the non-normal nature of the averaged ordinal fibrosis scores.

The linear regression does not directly estimate the dominance coefficient that interests us here. For instance, if there is no statistical interaction between genotypes (e.g., if *β_RB_* = 0), then fibrosis would be expected to equal the baseline *α* plus the independent effects of R and of B, yielding *y* = *α* + *β_R_* + *β_B_*, which is higher than either protein treatment alone (e.g., genetic overdominance). In contrast a genetically additive IGE should have fibrosis levels *α* + *β_B_* < *y* < *α* + *β_R_* which would require a negative *b_RB_* interaction effect. Therefore, we built a Bayesian linear model to directly estimate the dominance coefficient of the indirect genetic effects. We fit a model in which *y_i_*~*N*(*ŷ, σ*), and *ŷ* = *α* + *β_R_R* + *β_B_B* + *d*(*β_R_* – *β_B_*)*RB* to estimate the dominance coefficient *d* for Roselle and Boot lakes, and a similar analysis for Roselle and Cheney Lake coinjection. We extracted 1000 samples from the posterior distribution to estimate the mean and posterior predictive interval for each parameter. All analyses were conducted in R (R. Development Core Team 2022); data and code to reproduce analyses in this paper are archived for public access (https://doi.org/10.6084/m9.figshare.22083230.v1).

## EXPERIMENTAL RESULTS

### Experiment 1: Parasite indirect effects on host immune traits

Roselle Lake stickleback injected with tapeworm protein exhibited moderate fibrosis 10 days post-injection, consistent with prior results (Hund et al. 2022). The key insight from this experiment is that there were significant differences in fibrosis severity, depending on which tapeworm population was injected. Roselle Lake stickleback responded more strongly to Roselle Lake tapeworm protein, than to protein from lakes over 100 km away (Fig. 3). The among-population variation was statistically significant using either parametric or nonparametric tests (ANOVA: F_3,65_ = 5.37, P = 0.002); Kruskal-Wallis test χ^2^ = 10.74, df = 3, P = 0.0130). A Bayesian linear model estimated a larger effect for Roselle Lake cestode protein (β=0.67 [95% posterior predictive interval: 0.36-0.98]) than protein from other lakes (Boot Lake β= 0.01 [−0.30, 0.32], Gosling Lake β=0.09 [−0.22, 0.41]), using Cheney Lake as the baseline (Figure S2). We use these posterior probabilities as priors in analysis of experiment 2. These results confirm that protein from different parasite populations induce different levels of host fibrosis. We provisionally interpret this as a case of an indirect genetic effect (IGE). However, we acknowledge a key caveat: the proteins used here were derived from parasites dissected from wild-caught fish from these different lakes and may retain environmentally-induced differences (including differences induced by their original fish hosts).

**Figure 3.**
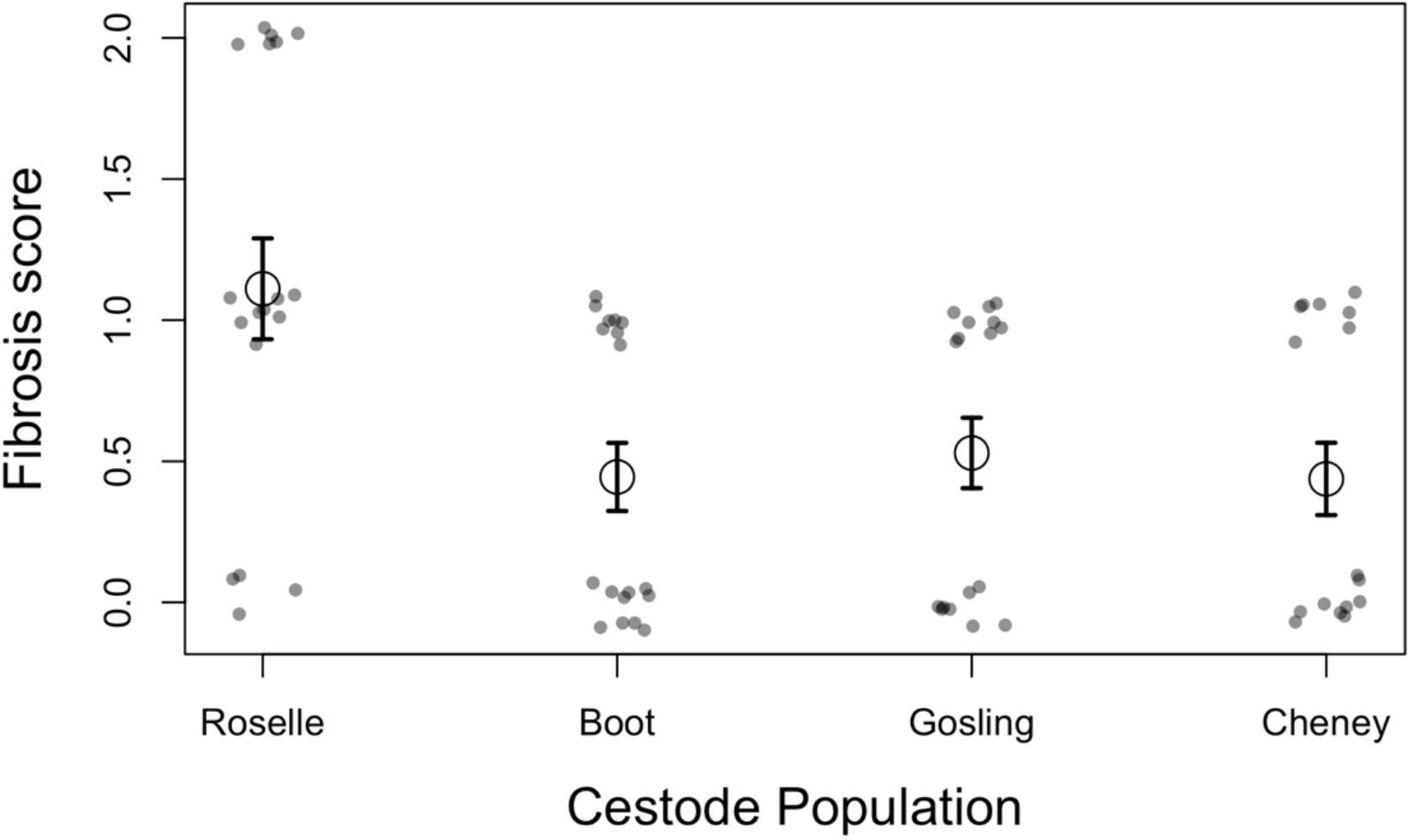
Fibrosis score (ordinal, jitter added to separate individual data points) as a function of the genotype of cestode protein injected into Roselle Lake stickleback. The mean of each treatment is denoted by a larger circle with ±1 standard error bars.

### Experiment 2: Overdominant indirect genetic effects during coinfection

We first compared whether fibrosis differed between coinjected fish receiving low versus high protein concentrations (substitutive versus additive treatments). We found no significant effect of concentration in either coinjection combination (Roselle + Boot: t = 1.524, p = 0.134; Roselle + Cheney: t = −1.394, p = 0.169). A subsequent experiment (Figure S3) subsequently confirmed that fibrosis is insensitive to a wide range of cestode protein concentrations. Consequently, for simplicity the following analyses present analyses that omit the effect of concentration, merging the additive and multiplicate treatments for a given combination of parasite proteins

The injected stickleback from Roselle Lake exhibited low levels of fibrosis when injected with saline (PBS controls, mean score = 1.2), or after injection with protein from the two geographically distant lakes (Boot: mean = 1.372, t = 0.650, p = 0.516; Cheney: mean = 1.407, t = 0.799, p = 0.425). However, fish injected with Roselle Lake cestode protein experienced somewhat higher fibrosis than the control fish (mean = 1.630, t = 1.654, p = 0.099), consistent with Experiment 1. Both coinjection groups (each containing Roselle protein) exhibited significantly elevated fibrosis relative to the control (Roselle + Boot: mean = 1.967, t = 3.330, p = 0.001; Roselle + Cheney: mean = 1.694, t = 2.186, p = 0.030, Table 3, Figure 4). An ANOVA confirmed that the presence of Roselle Lake protein (whether alone or in combination) significantly increased fibrosis (Table 1). We found no statistical interaction between worm protein genotypes, suggesting they have a statistically additive effect. However, from a biological standpoint this statistically additive effect implies genetic overdominance: greater fibrosis for the coinjected fish than for fish receiving either genotype’s protein alone (Fig. 4).

**Figure 4.**
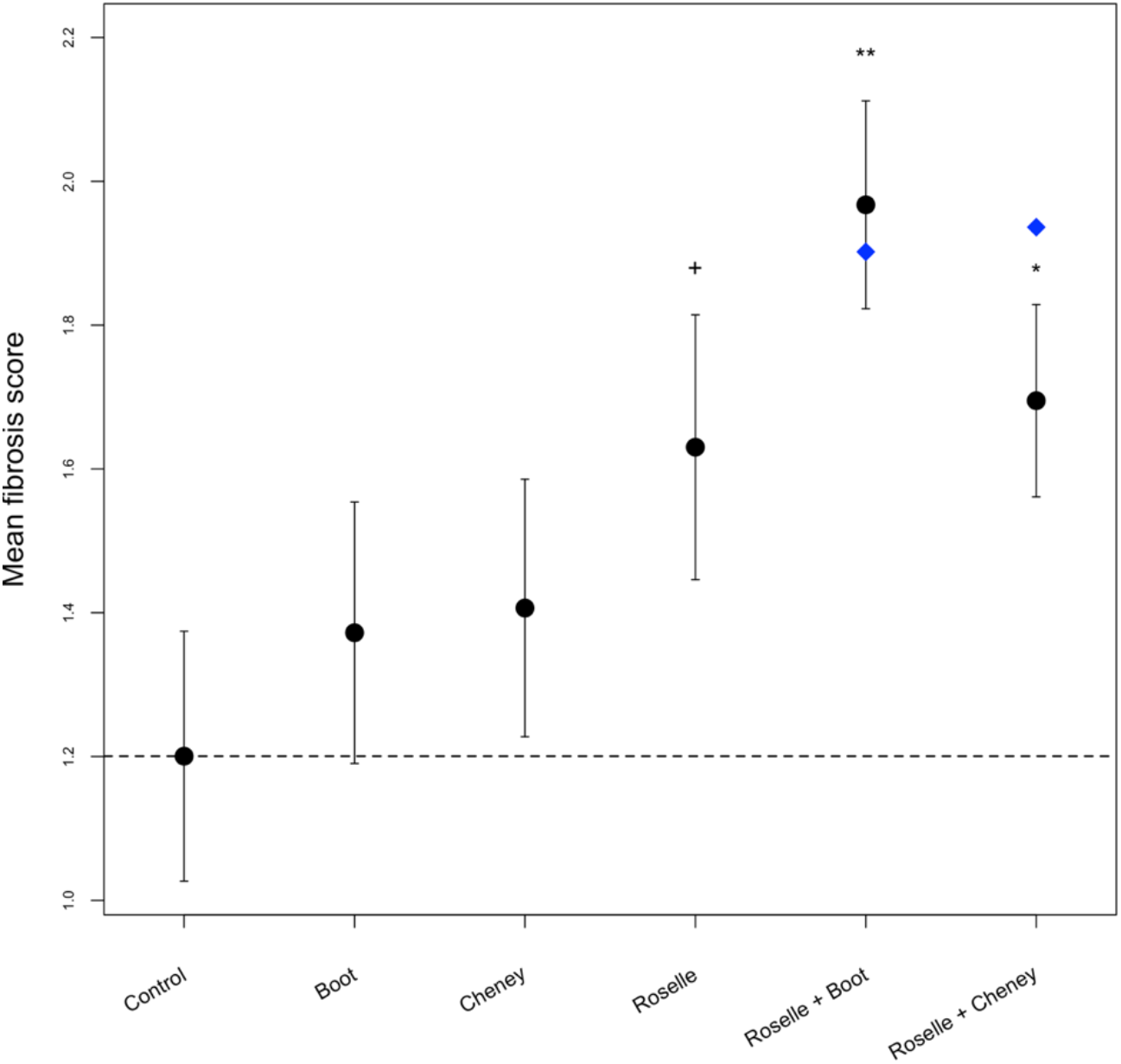
Mean fibrosis score of saline-injected stickleback (dashed line for reference), and fish injected with single or combined tapeworm proteins (merging additive and substitutive concentrations). We present the means (solid circles) with ±1 standard error bars. Asterisks denote significant differences from the control (+ P<0.1; * P<0.05; ** P<0.01). The blue diamond indicates the expected value under a strictly additive statistical model (not to be confused with an additive genetic model, where the point should fall between the values for the two protein genotypes injected separately). Figure S4 presents the raw data values.

**Table 1:**
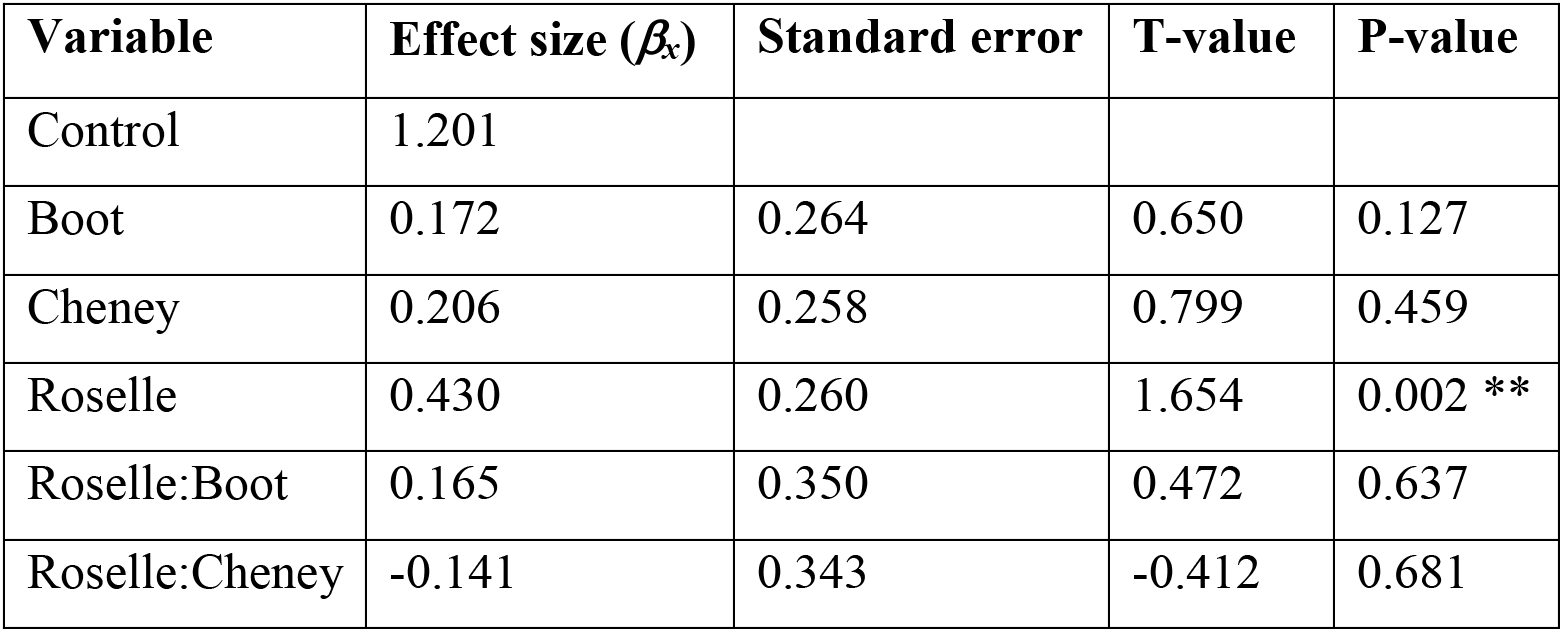
Effect sizes, standard error of effect, t-values, and p-values for all possible predicting factors of fibrosis levels (not including sex-or mass-dependent interactions). Significance of effect size is indicated as follows: *** indicates p < 0.001, ** indicates p < 0.01, * indicates p < 0.05, and + indicates p < 0.1.

| Variable | Effect size ($\beta_x$) | Standard error | T-value | P-value |
| --- | --- | --- | --- | --- |
| Control | 1.201 |  |  |  |
| Boot | 0.172 | 0.264 | 0.650 | 0.127 |
| Cheney | 0.206 | 0.258 | 0.799 | 0.459 |
| Roselle | 0.430 | 0.260 | 1.654 | 0.002 ** |
| Roselle:Boot | 0.165 | 0.350 | 0.472 | 0.637 |
| Roselle:Cheney | -0.141 | 0.343 | -0.412 | 0.681 |

We estimated the dominance coefficient using a Bayesian linear model using priors derived from Experiment 1. Focusing first on the Roselle + Boot Lake combination (Figures S5–6), we confirmed that Roselle Lake fish respond with stronger fibrosis when injected with Roselle Lake tapeworm protein, compared to controls (β_Ros_ = 0.41, predictive interval [0.010, 0.72]). In contrast Roselle Lake fish exhibited negligible response to Boot Lake protein (β_Boot_ = 0.02, predictive interval [−0.27, 0.31]). The mean of the posterior distribution of the dominance coefficient was greater than 1.0 (implying overdominance), though the predictive interval was broad (β_D_ = 1.64, [0.26, 3.01]). Similar results were obtained for Roselle and Cheney Lake combinations (Figures S7–8): the Cheney Lake protein induced no more fibrosis than the saline controls (β_Cheney_ = 0.10, predictive interval [−0.20, 0.39]), whereas Roselle protein did (β_Ros_ = 0.43, predictive interval [0.11, 0.75]). The estimated dominance coefficient was again greater than one (β_D_ = 1.25, [−0.11, 2.61]). Although the dominance coefficients for both analyses yielded broad posterior predictive intervals, in both cases the best estimate was greater than 1.0, indicating that the Roselle Lake tapeworms’ pro-fibrosis IGE is at a minimum dominant over non-fibrotic Boot or Cheney Lake proteins. More likely Roselle and Boot combinations yield an overdominant effect, with a stronger immune response to the coinjected combination, than to Roselle alone. If we instead calculate the dominance coefficient from the frequentist linear model coefficient estimates, we infer that Roselle+Boot yield an overdominant effect (*d* = 2.306) while Roselle + Cheney tend in the same direction but less strongly (*d* = 1.289).

## DISCUSSION

In the coevolutionary arms race between hosts and parasites, parasites can gain an edge over their host by evading detection by the host’s immune receptors, or by secreting proteins that actively suppress host immunity (Schmid-Hempel 2008). Either strategy may reduce the host immune response thereby increasing parasite fitness, so at first glance the distinction may seem inconsequential. But when parasites with different strategies coinfect a single host, the distinction can matter greatly. Coinfection by both camouflaged and recognized parasite genotypes should enable host recognition. The more immune-stimulating parasite will have a dominant indirect genetic effect, simultaneously reducing the fitness of otherwise camouflaged genotype. In contrast, coinfection by suppressing and non-suppressing genotypes should still suppress the host response, rescuing the fitness of the latter genotype.

As we demonstrated with the model presented here, the dominance of an indirect genetic effect will fundamentally alter the parasite’s evolutionary dynamics. In particular, the relative fitness of an immigrant parasite genotype is contingent on (1) the probability it ends up in a coinfected host (almost always with a resident parasite genotype), and (2) whether its immune evasion is maintained or undermined by that coinfection. If resident parasites stimulate a stronger host immune response, and this IGE is dominant, then immune-evasive immigrants will have difficulty invading a high-prevalence population. Conversely, when residents are rare, immigrants maintain their immune-evasive advantages. Thus, the effective rate of gene flow between parasite populations, and thus the long-term trajectory of parasite population divergence (or, introgression) depends on an interaction between IGE dominance and resident infection rates.

Indirect genetic effects have been studied for some time (Baud et al. 2022), particularly in the context of social behavior (Santostefano et al. 2017). More recently IGEs have been applied to study coevolution between hosts and parasites (De Lisle et al. 2022). However, to the best of our knowledge the issue of IGE dominance has not previously been considered. Nor do we have much empirical data to evaluate IGE dominance. Our antigen injection experiment provides one case study to confirm that IGEs occur in host-parasite interactions, and exhibit a form of dominance. Cestode antigen injection induces a host immune response in stickleback (fibrosis), consistent with other studies that reported *S.solidus* manipulation of host immune traits (Scharsack et al. 2004, 2007b, 2013; Berger et al. 2021). Crucially, we demonstrated here that antigens from different parasite populations induce correspondingly different levels of fibrosis, in a shared host genetic background (Roselle Lake fish). In both experiment 1 and 2, Roselle Lake stickleback responded more strongly to Roselle Lake cestode protein, than to antigens from allopatric parasite populations. Thus, the different host fibrosis traits likely represent a parasite IGE. A key caveat is that the injected parasite proteins were obtained from wild-caught parasites that may retain divergent environmental effects on their proteomes. It would be helpful to follow this study with a multi-generation common-garden rearing design to obtain a more robust inference about parasite genetic differences. That said, there are genetic differences between the parasite populations studied here, including loci where genetic divergence correlates with host fibrosis traits (Shim et al. 2022b), despite otherwise low genome-wide genetic differentiation (Shim et al. 2022a).

Having documented likely parasite IGEs affecting a host immune trait (fibrosis), we were then able to leverage the convenience of the injection protocol to evaluate the consequences of coinfection (e.g., coinjection). To our surprise, the coinjected fish exhibited a consistently stronger response than either genotype alone, even with the substitutive design that kept total antigen concentration constant. Thus, the fibrosis-inducing genotype is at a minimum dominant; our best estimate is that the indirect genetic effects are overdominant (for both protein combinations). The immunological molecular mechanisms of this overdominance are not currently known, nor do we know what parasite antigens are recognized by the host, nor the mechanisms of host recognition. However, we can infer that the different indirect genetic effects are more consistent with a recognition-success model (Figure 1A), which should generate a dominant IGE, as opposed to a parasite immune-suppression model (Figure 1B) where we expect the IGE to be recessive.

Our model suggests that this overdominance will tend to inhibit gene flow from foreign parasite populations, especially when resident infection rates are high (e.g., coinfection is common). Fish-eating birds are the definitive host of *S. solidus* (Clarke 1954), which they acquire by eating infected stickleback. Because birds are so vagile, they readily distribute tapeworm eggs across a variety of populations. Indeed, a population genetic survey of a dozen lakes on Vancouver Island identified individual parasites that are likely first generation or F1 immigrants (Shim et al. 2022b), and F_ST_ between lakes is generally negligible. Isolation by distance exists but is weak even at a scale of hundreds of kilometers. Thus, gene flow is likely to be a substantial force in *S.solidus* evolution. The dominance of parasite IGEs should therefore be an important factor regulating rates of population genetic divergence, local adaptation, and coevolution (Gandon and Michalakis 2002).

More generally, we suggest that the dominance of indirect genetic effects deserves more extensive attention in a variety of systems. This will be relevant any time two conditions are met: (1) indirect genetic effects exist, and (2) more than one individual of the causal species is exerting influence on the same recipient. This is likely to be a widespread situation in many coevolutionary interactions, including host-parasite interactions (De Lisle et al. 2022). For instance, helminth-induced modulation of the human immune system can interfere with vaccine efficacy (Labeaud et al. 2009; Moreau and Chauvin 2010; Wait et al. 2020), and helminth coinfection rates are high in many human populations (Hoetz et al. 2008). There are many open avenues for research that incorporates IGE dominance in host-parasite coevolutionary theory and epidemiology.

## Acknowledgements

We thank Sarah Tsuruo for assistance with the injection experiments, and Emma Choi and Maria Rodgers for comments on initial drafts of the paper. This research was supported by NIH grant 1R01AI123659-01A1 to DIB, and by the University of Connecticut.

## Author contributions

Experiment 1 was designed by DIB and CP, and conducted by CP. Experiment 2 was designed and conducted by SA, AR, LS, and DIB as part of SA’s undergraduate thesis. AP contributed the experiment in Fig. S3. DIB conducted the data analyses, model conception and analysis. The manuscript was written by DIB and SA with feedback from all authors. The authors have no competing interests to declare.

## Data Accessibility

All data and original R code needed to reproduce the results reported in this paper are publicly available on FigShare (https://doi.org/10.6084/m9.figshare.22083230.v1).

---

**Table S1:**
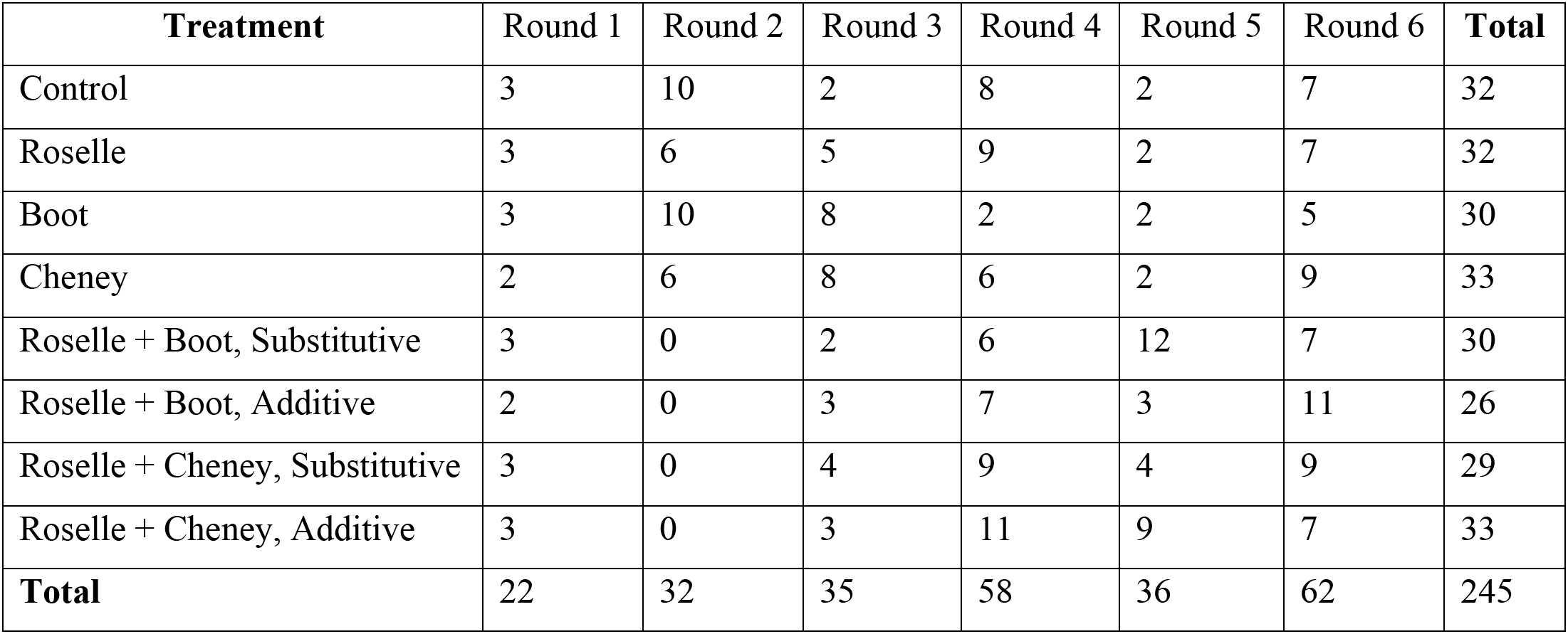
Number of fish that we factored into our analyses, after necessary discards due to a) deaths and b) sample size discrepancies identified at the time of dissection.

**Figure S1.**
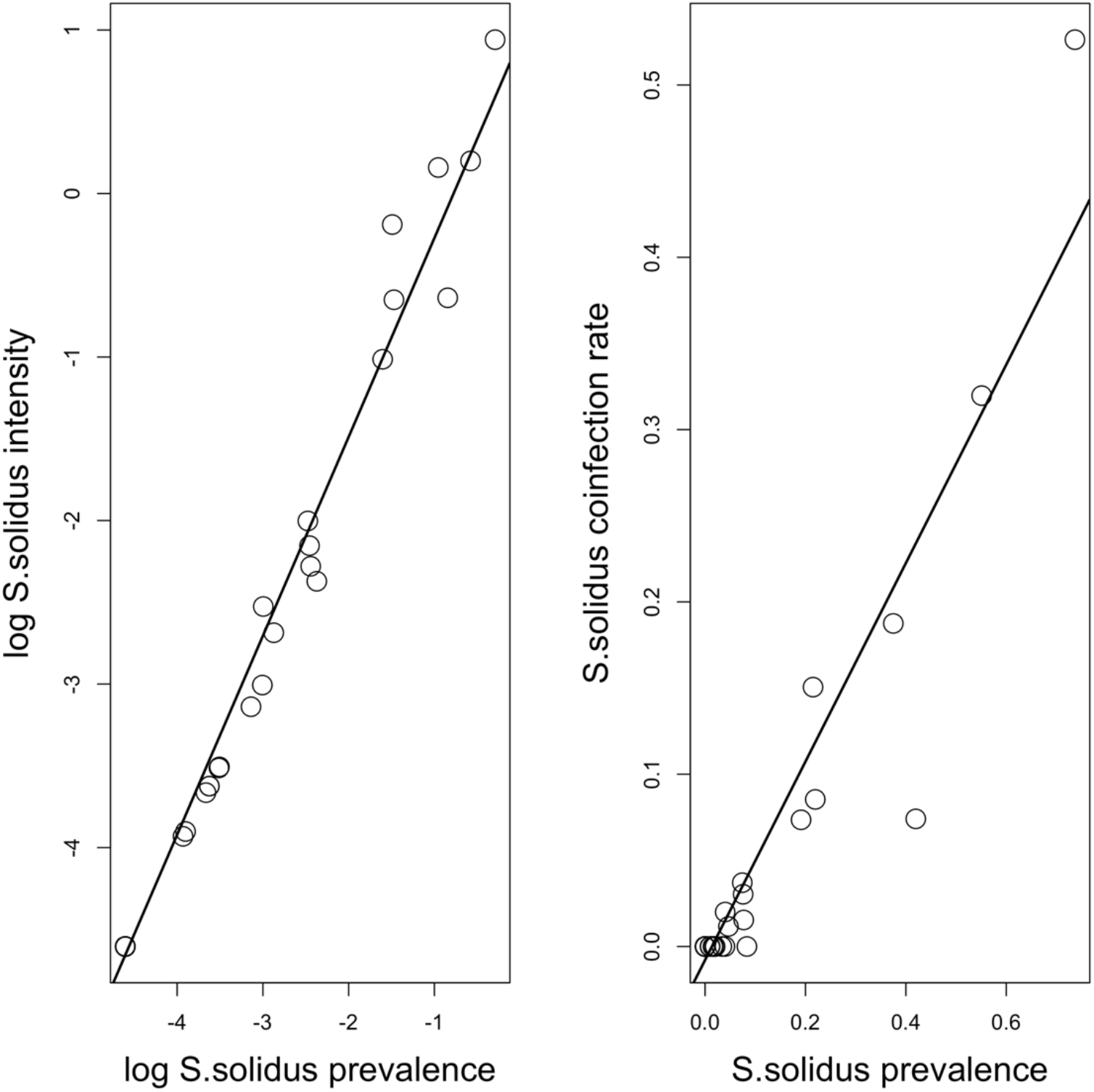
Re-analysis of previously published data on *S.solidus* infection rates in lake populations of stickleback from Vancouver Island (Bolnick et al. 2020). Up to 200 stickleback were sampled from each of 46 populations in 2009, and dissected to determine the number of *S.solidus.* We calculated prevalence as the proportion of fish with infections present, intensity as the mean number of *S.solidus* per individual (including uninfected cases), and coinfection rate as the proportion of fish with more than one infection. Lines represent the best fit regression (t = 53.8, t = 18.8 respectively, both P < 0.0001).

**Figure S2.**
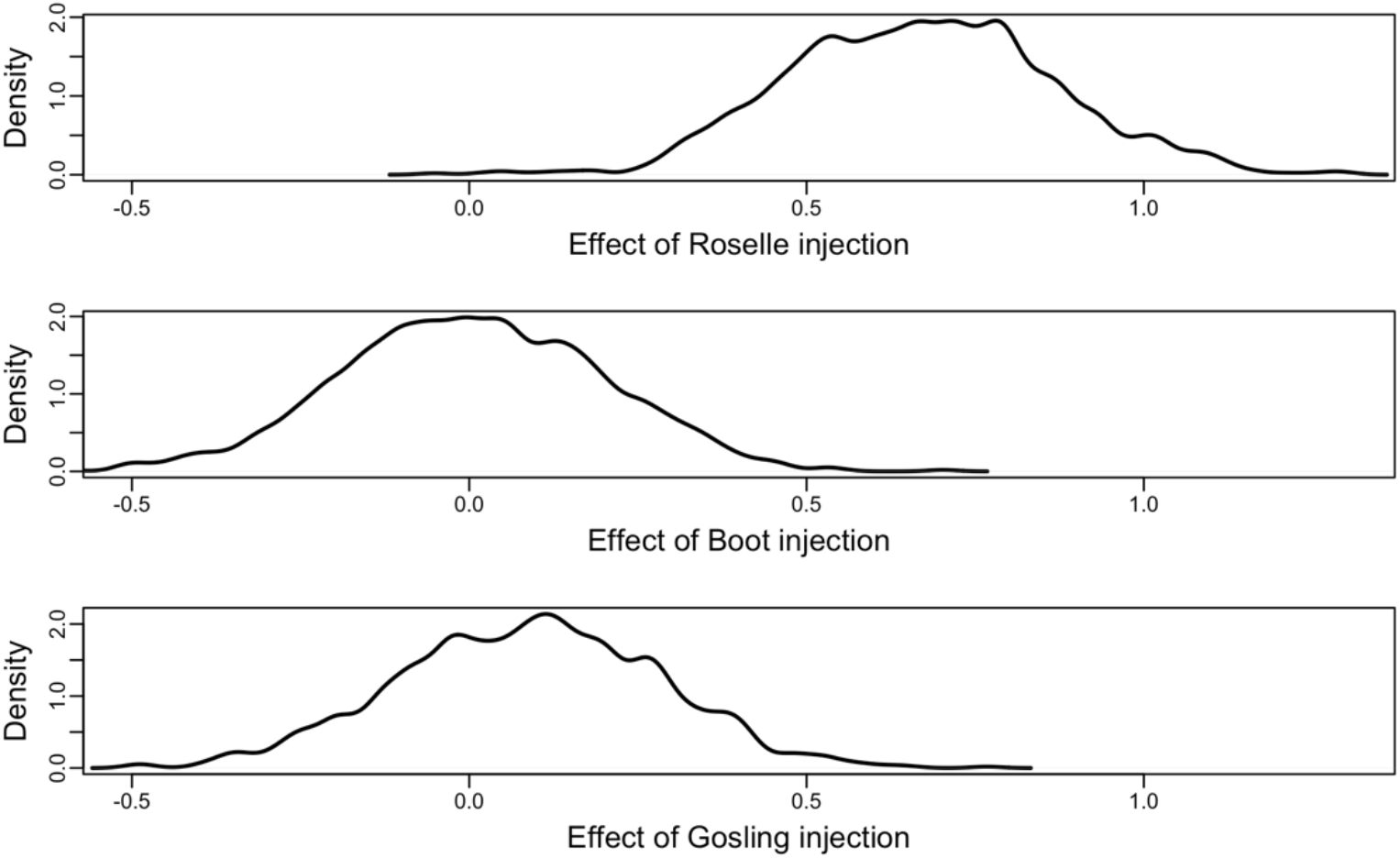
Posterior distribution of Experiment 1 estimates of effect sizes β_R_, β_B_, and β_G_ relative to the most distant Cheney Lake used as a baseline (a).

**Figure S3.**
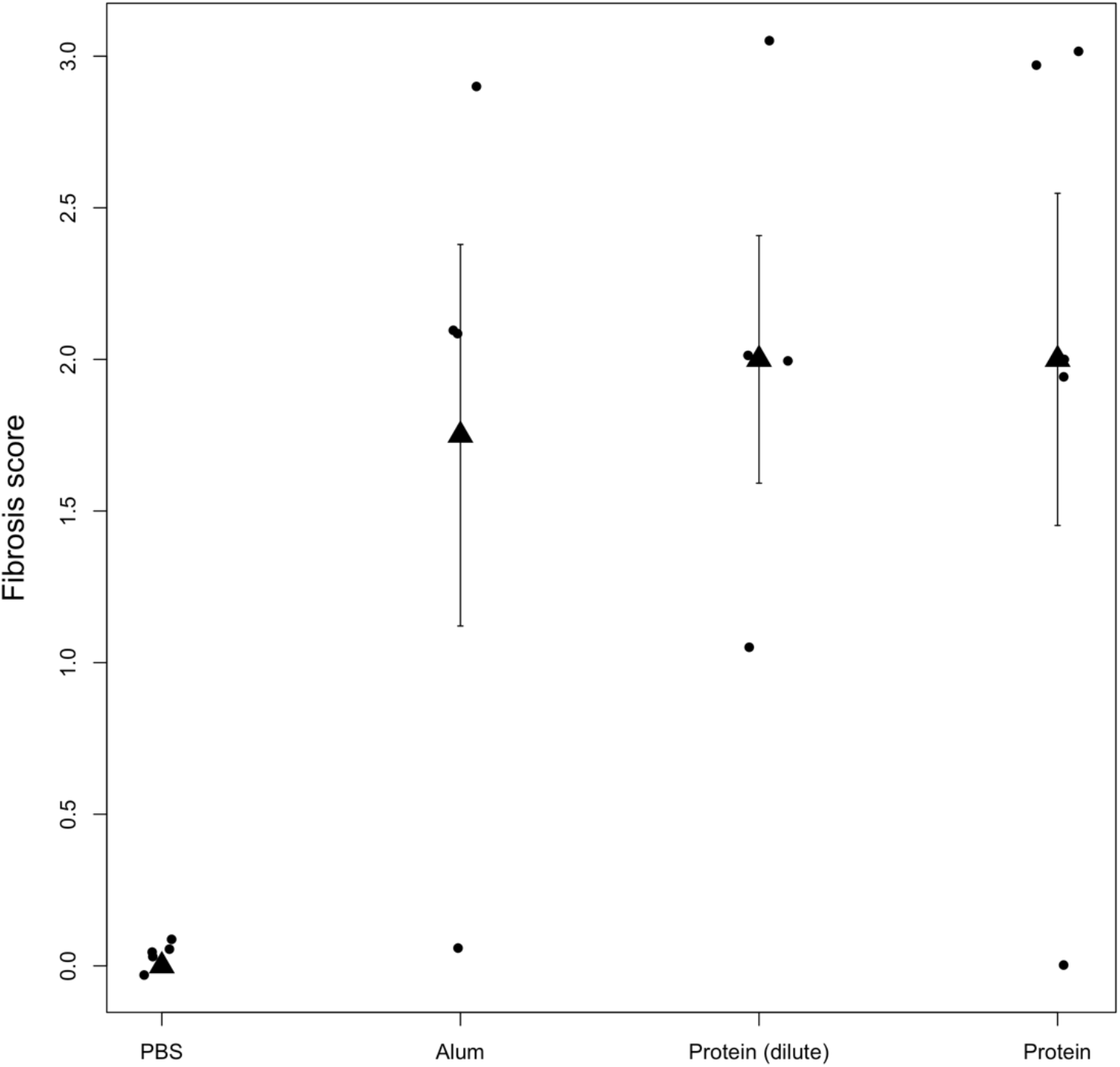
A follow-up experiment re-evaluated the lack of a protein concentration effect, using a three order-of-magnitude dilution rather than a mere doubling. We injected lab-raised stickleback from Boot Lake (which also exhibit strong fibrosis) with one of four treatments: a negative control treatment (20 μL of of 0.9x PBS), a positive control treatment (alum), and two protein injections: the full concentration 20 μL of 1 mg/ml protein from Boot Lake cestodes in 0.9x PBS, or a 1/1000 dilution. A linear model confirmed significant differences between the three treatments and the negative control (alum t = 2.73, P = 0.0162; protein t = 3.12, P = 0.0075; diluted protein t = 3.31, P = 0.0051).

**Figure S4.**
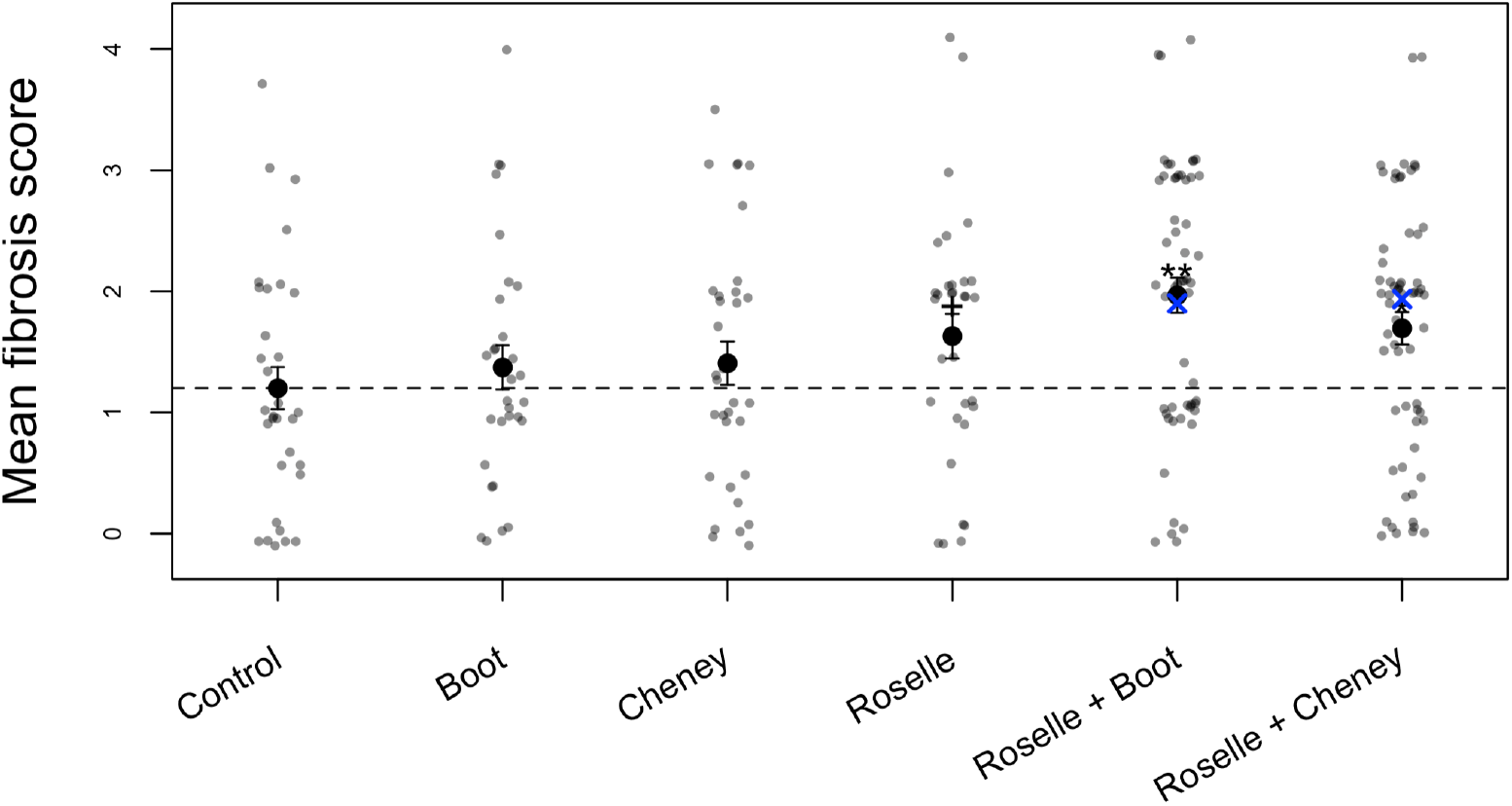
Raw data corresponding to Figure 4, from Experiment 2.

**Figure S5.**
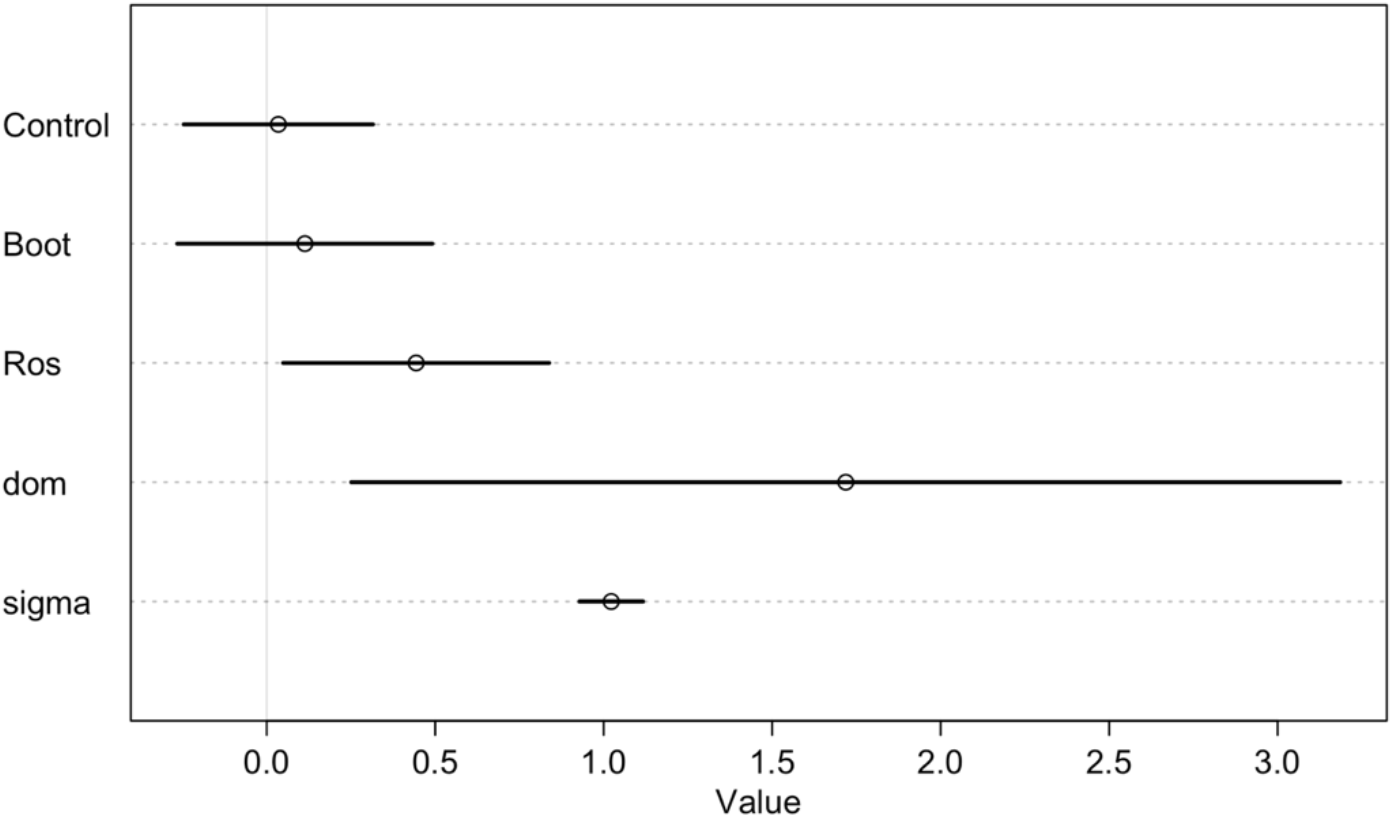
95% posterior predictive intervals from the Bayesian analysis of Experiment 2 injections of Boot Lake protein, Roselle Lake protein, or coinjection (saline control mean α, and effects β_B_, β_R_, and dominance coefficient d), with overall sampling standard deviation σ.

**Figure S6.**
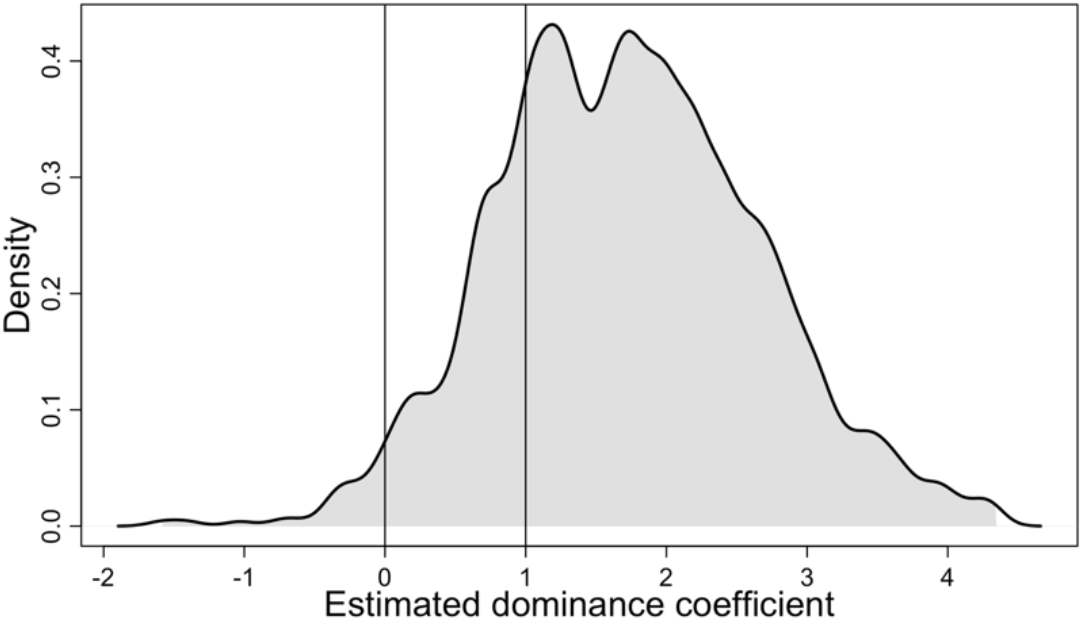
Histogram of 1000 samples from the posterior distribution of estimates of the Roselle and Boot Lake dominance coefficient d. For values greater than 1.0, fish respond with stronger fibrosis to the combined injection, than to either injection alone. Values of 0 imply the lower-fibrosis Boot Lake dominates (consistent with an immune suppression model). Values of 1 imply the higher-fibrosis Roselle Lake dominates (consistent with a parasite-detection model). Values between 0 and 1 would imply partially dominant or additive effects.

**Figure S7.**
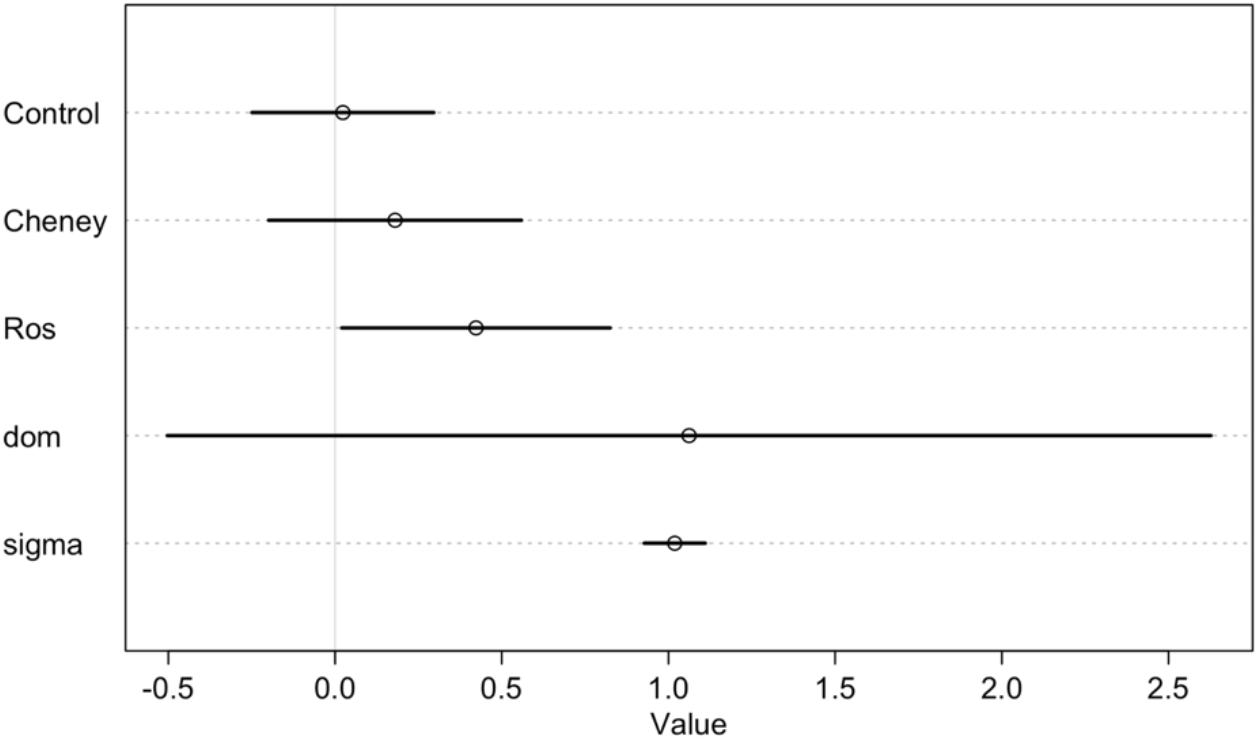
95% posterior predictive intervals from the Bayesian analysis of Experiment 2 injections of Cheney Lake protein, Roselle lake protein, or coinjection (saline control mean a, and effects β_C_, β_R_, and dominance coefficient d), with overall sampling standard deviation σ.

**Figure S8.**
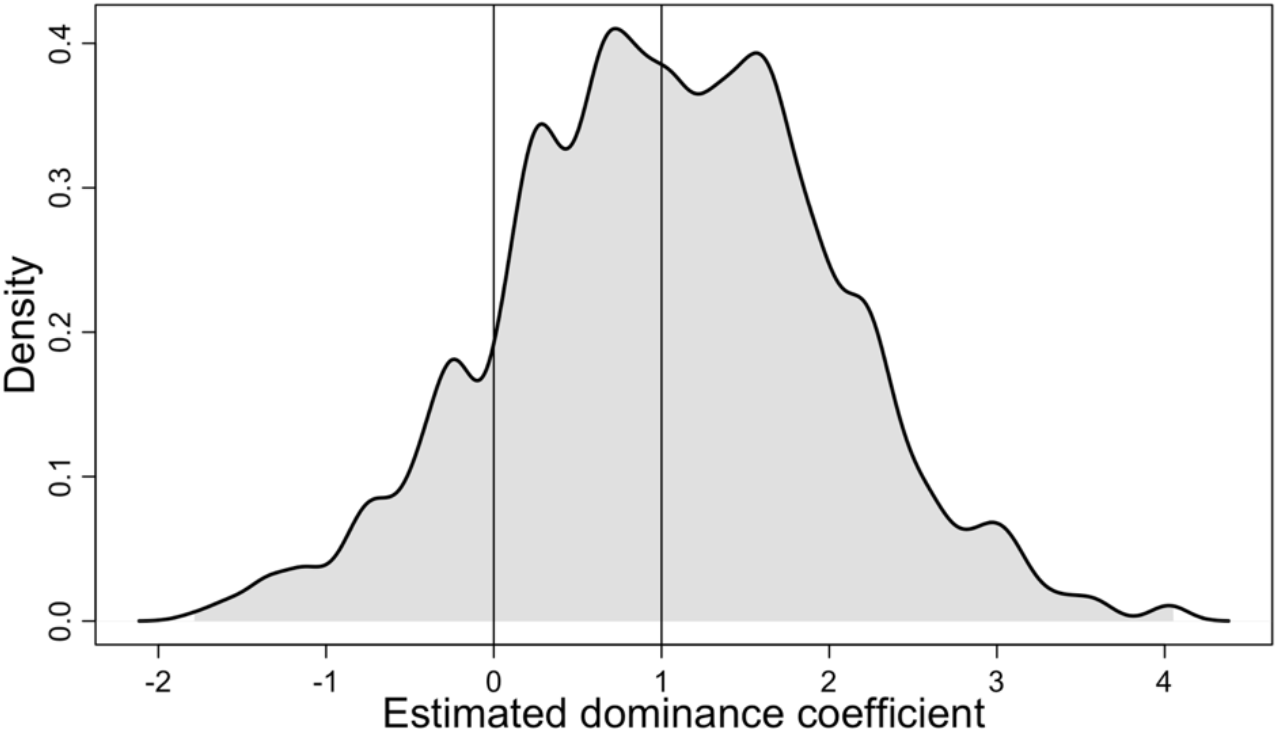
Histogram of 1000 samples from the posterior distribution of estimates of the Roselle and Cheney Lake dominance coefficient d. For values greater than 1.0, fish respond with stronger fibrosis to the combined injection, than to either injection alone. Values of 0 imply the lower-fibrosis Cheney Lake dominates (consistent with an immune suppression model). Values of 1 imply the higher-fibrosis Roselle Lake dominates (consistent with a parasite-detection model). Values between 0 and 1 would imply partially dominant or additive effects.

