## Supplemental Tables and Figures for "The dominance of coinfecting parasites’ indirect effects on host traits"

**Table S1:** Number of fish that we factored into our analyses, after necessary discards due to a) deaths and b) sample size discrepancies identified at the time of dissection.

| **Treatment** | Round 1 | Round 2 | Round 3 | Round 4 | Round 5 | Round 6 | **Total** |
|---|---|---|---|---|---|---|---|
| Control | 3 | 10 | 2 | 8 | 2 | 7 | 32 |
| Roselle | 3 | 6 | 5 | 9 | 2 | 7 | 32 |
| Boot | 3 | 10 | 8 | 2 | 2 | 5 | 30 |
| Cheney | 2 | 6 | 8 | 6 | 2 | 9 | 33 |
| Roselle + Boot, Substitutive | 3 | 0 | 2 | 6 | 12 | 7 | 30 |
| Roselle + Boot, Additive | 2 | 0 | 3 | 7 | 3 | 11 | 26 |
| Roselle + Cheney, Substitutive | 3 | 0 | 4 | 9 | 4 | 9 | 29 |
| Roselle + Cheney, Additive | 3 | 0 | 3 | 11 | 9 | 7 | 33 |
| **Total** | 22 | 32 | 35 | 58 | 36 | 62 | 245 |

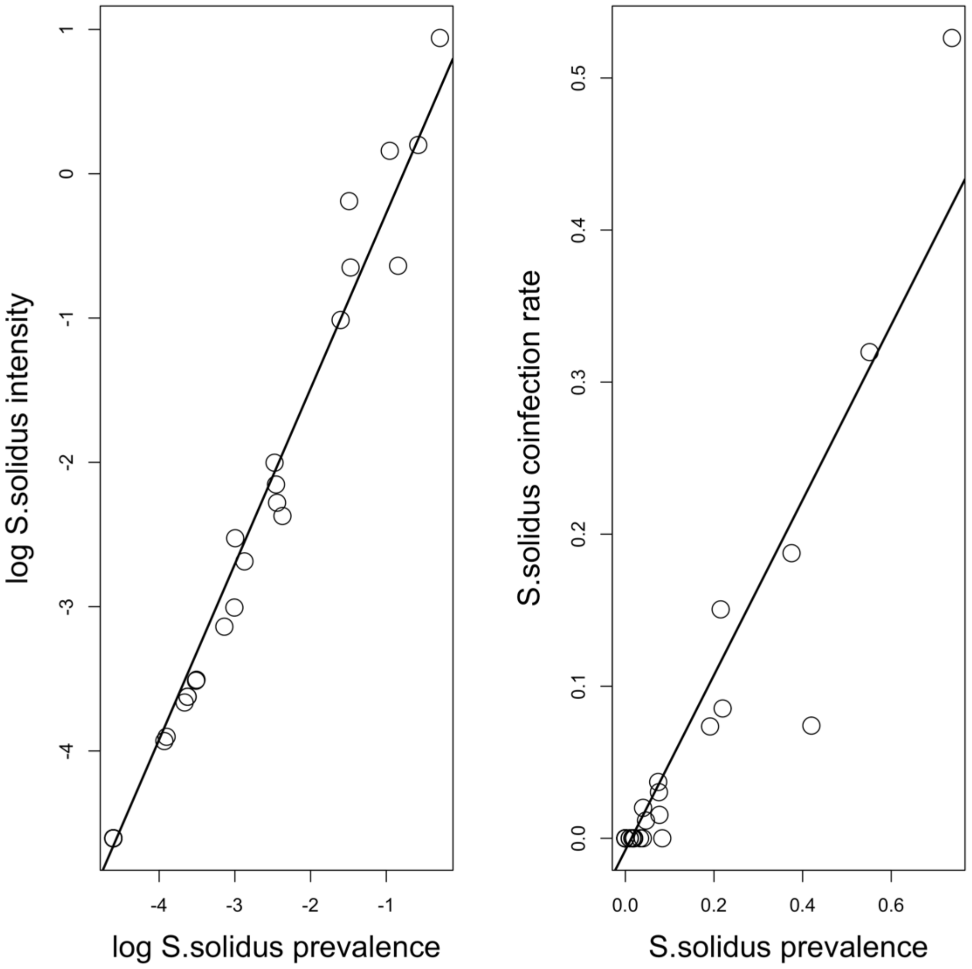


**Figure S1.** Re-analysis of previously published data on *S.solidus* infection rates in lake populations of stickleback from Vancouver Island (Bolnick et al. 2020). Up to 200 stickleback were sampled from each of 46 populations in 2009, and dissected to determine the number of *S.solidus*. We calculated prevalence as the proportion of fish with infections present, intensity as the mean number of *S.solidus* per individual (including uninfected cases), and coinfection rate as the proportion of fish with more than one infection. Lines represent the best fit regression ($t = 53.8$, $t = 18.8$ respectively, both $P < 0.0001$).

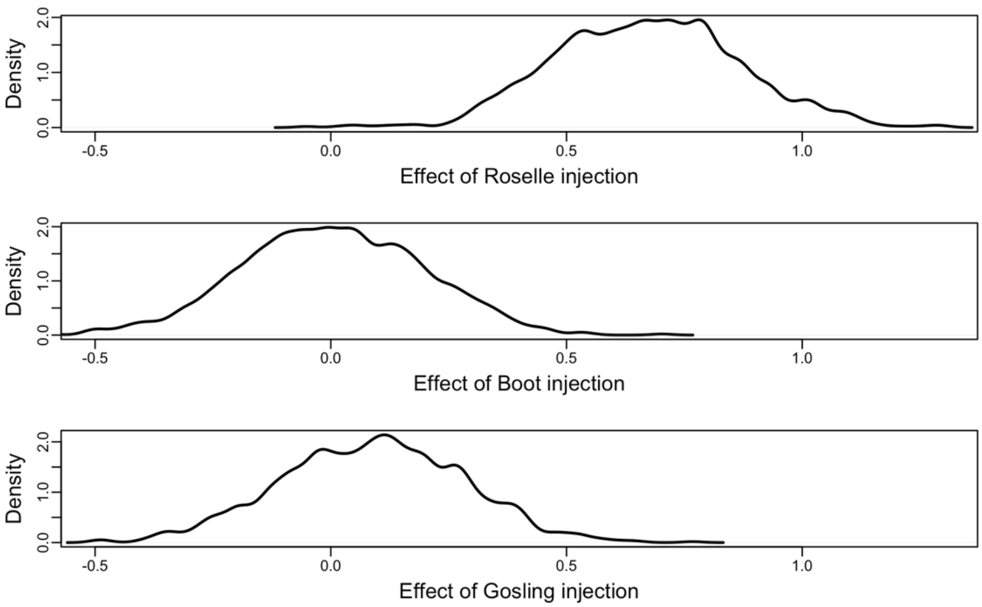


**Figure S2.** Posterior distribution of Experiment 1 estimates of effect sizes $\beta_R$, $\beta_B$, and $\beta_G$ relative to the most distant Cheney Lake used as a baseline ($\alpha$).

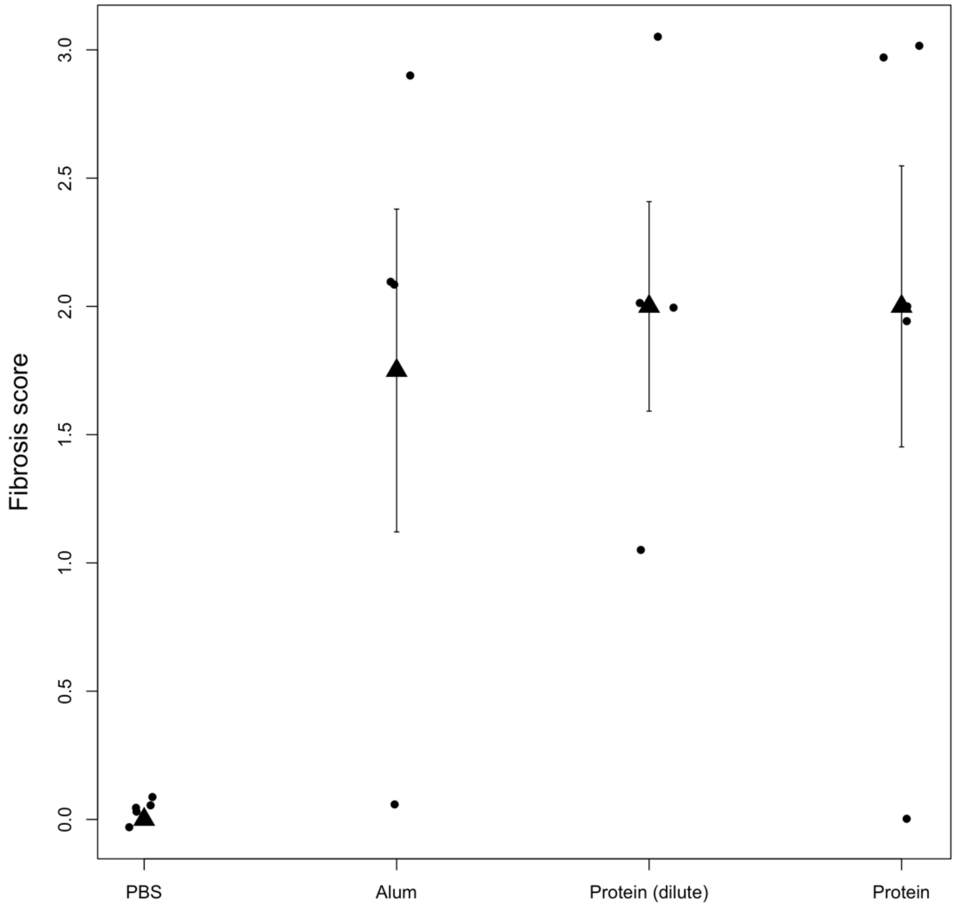


**Figure S3:** A follow-up experiment re-evaluated the lack of a protein concentration effect, using a three order-of-magnitude dilution rather than a mere doubling. We injected lab-raised stickleback from Boot Lake (which also exhibit strong fibrosis) with one of four treatments: a negative control treatment (20 μL of of 0.9x PBS), a positive control treatment (alum), and two protein injections: the full concentration 20 μL of 1 mg/ml protein from Boot Lake cestodes in 0.9x PBS, or a 1/1000 dilution. A linear model confirmed significant differences between the three treatments and the negative control (alum $t = 2.73$, $P = 0.0162$; protein $t = 3.12$, $P = 0.0075$; diluted protein $t = 3.31$, $P = 0.0051$).

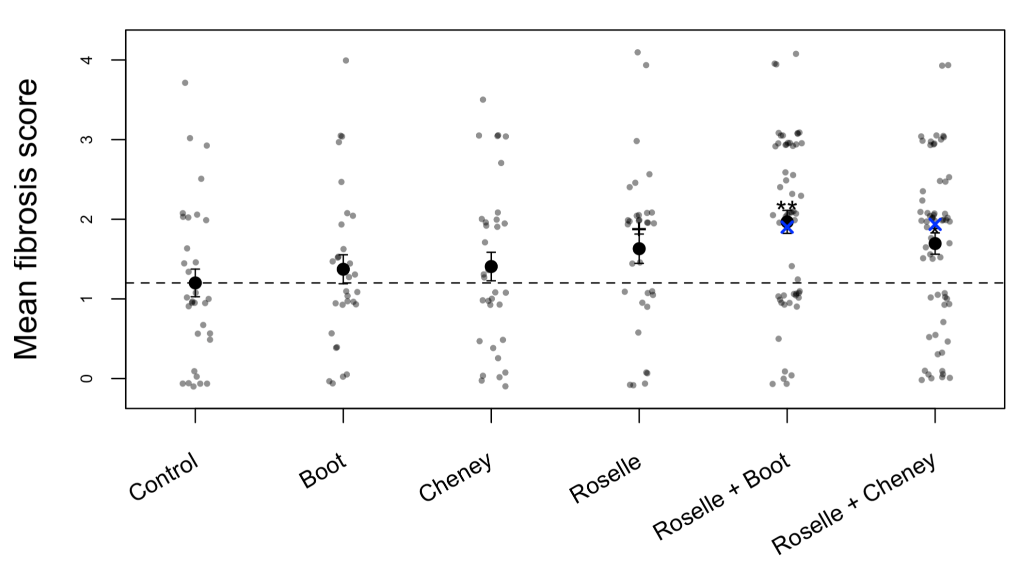


**Figure S4**. Raw data corresponding to Figure 4, from Experiment 2.

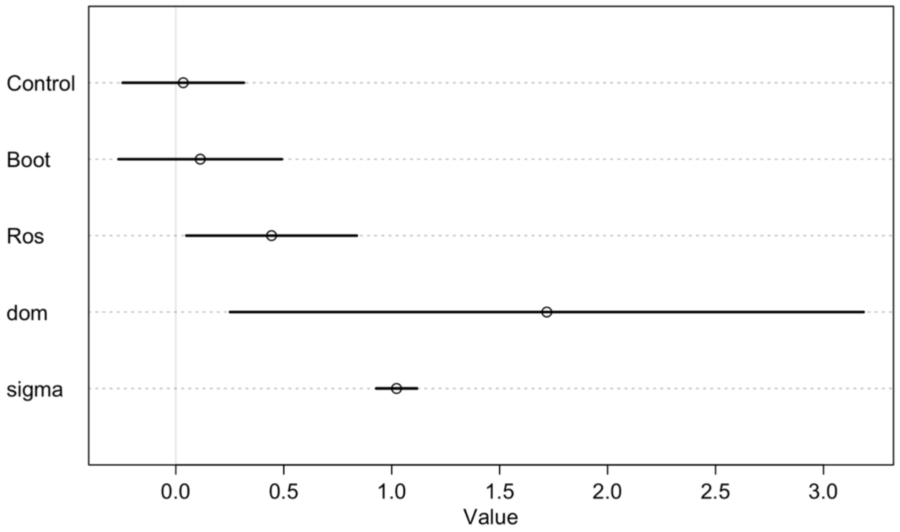


**Figure S5.** 95% posterior predictive intervals from the Bayesian analysis of Experiment 2 injections of Boot Lake protein, Roselle Lake protein, or coinjection (saline control mean $\alpha$, and effects $\beta_B$, $\beta_R$, and dominance coefficient d), with overall sampling standard deviation $\sigma$.

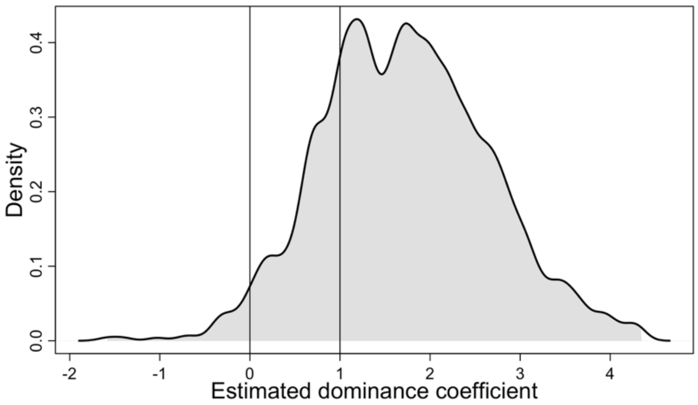


**Figure S6.** Histogram of 1000 samples from the posterior distribution of estimates of the Roselle and Boot Lake dominance coefficient d.  For values greater than 1.0, fish respond with stronger fibrosis to the combined injection, than to either injection alone. Values of 0 imply the lower-fibrosis Boot Lake dominates (consistent with an immune suppression model). Values of 1 imply the higher-fibrosis Roselle Lake dominates (consistent with a parasite-detection model). Values between 0 and 1 would imply partially dominant or additive effects.

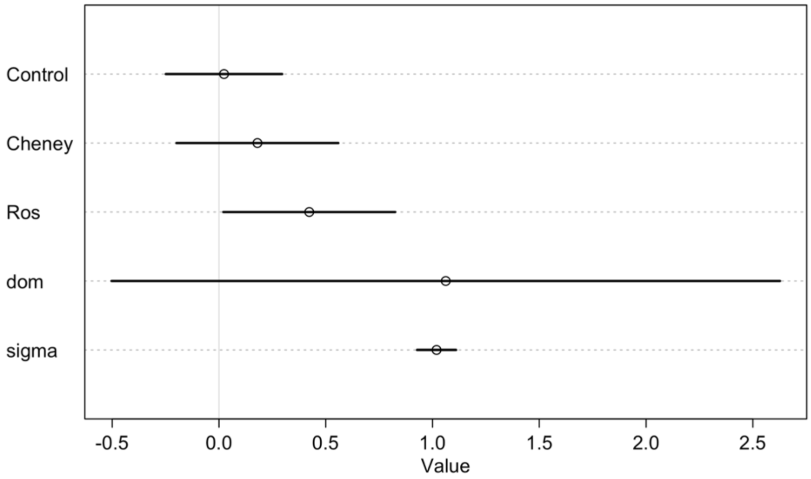


**Figure S7.** 95% posterior predictive intervals from the Bayesian analysis of Experiment 2 injections of Cheney Lake protein, Roselle lake protein, or coinjection (saline control mean $\alpha$, and effects $\beta_C$, $\beta_R$, and dominance coefficient d), with overall sampling standard deviation $\sigma$.

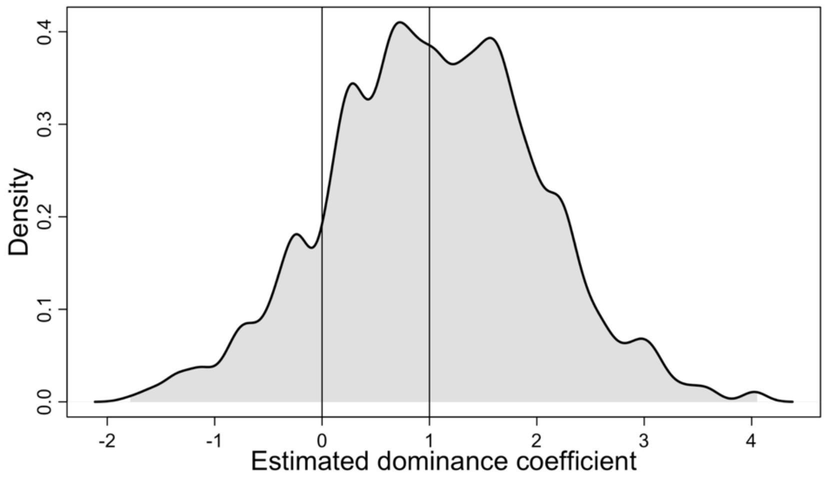


**Figure S8.** Histogram of 1000 samples from the posterior distribution of estimates of the Roselle and Cheney Lake dominance coefficient d. For values greater than 1.0, fish respond with stronger fibrosis to the combined injection, than to either injection alone. Values of 0 imply the lower-fibrosis Cheney Lake dominates (consistent with an immune suppression model). Values of 1 imply the higher-fibrosis Roselle Lake dominates (consistent with a parasite-detection model). Values between 0 and 1 would imply partially dominant or additive effects.
